# Transcriptional activity mediated by β-CATENIN and TCF/LEF family members is completely dispensable for survival of multiple human colorectal cancer cell lines

**DOI:** 10.1101/2022.12.05.519142

**Authors:** Janna Fröhlich, Katja Rose, Andreas Hecht

## Abstract

Unrestrained transcriptional activity of β-CATENIN and its binding partner TCF7L2 frequently underlies colorectal tumor initiation and is considered an obligatory oncogenic driver throughout intestinal carcinogenesis. Yet, the *TCF7L2* gene carries inactivating mutations in about 10 % of colorectal tumors and is non-essential in colorectal cancer (CRC) cell lines. To determine whether CRC cells acquire TCF7L2-independence through cancer-specific compensation by other T-cell factor (TCF)/lymphoid enhancer-binding factor (LEF) family members, or rather lose addiction to β-CATENIN/TCF7L2-driven gene expression altogether, we generated multiple CRC cell lines entirely negative for TCF/LEF or β-CATENIN expression. Viability of these cells demonstrates complete β-CATENIN- and TCF/LEF-independence, albeit one β-CATENIN-deficient cell line eventually became senescent. Absence of TCF/LEF proteins and β-CATENIN consistently impaired CRC cell proliferation, reminiscent of mitogenic effects of WNT/β-CATENIN signaling in the healthy intestine. Despite this common phenotype, β-CATENIN-deficient cells exhibited highly cell-line-specific gene expression changes with little overlap between β-CATENIN- and TCF7L2-dependent transcriptomes. Apparently, β-CATENIN and TCF7L2 control sizeable fractions of their target genes independently from each other. The observed divergence of β-CATENIN and TCF7L2 transcriptional programs, and the finding that neither β-CATENIN nor TCF/LEF activity is strictly required for CRC cell survival has important implications when evaluating these factors as potential drug targets.

## Introduction

The WNT/β-CATENIN pathway is an evolutionarily conserved metazoan signal transduction cascade with pivotal functions throughout embryogenesis and in post-natal tissue homeostasis. Central to this pathway is the so-called destruction complex, a multifactorial assembly comprising among others the tumor suppressors AXIN and APC, CASEIN KINASE1α (CK1α), and GLYCOGEN SYNTHASE KINASE 3 (GSK3). Within the microcompartment of the destruction complex, CK1α and GSK3 sequentially phosphorylate β-CATENIN and thereby prime it for proteasomal degradation, thus keeping the pathway in the off state [1]. Binding of WNT growth factors to FRIZZLED and LRP cell surface receptors through mechanisms that are still not fully understood, inhibits the destruction complex. This results in β-CATENIN stabilization and translocation into the nucleus where it functions as cofactor of members of the TCF/LEF family of sequence specific DNA-binding proteins [2],[3]. Complexes formed by β-CATENIN and TCF/LEF proteins together with additional regulatory factors in turn elicit cell-type-specific and ubiquitous transcriptional responses [2],[3], which represent the main output of the WNT/β-CATENIN pathway.

Beginning in late stage mouse embryonic development, WNT/β-CATENIN signaling is absolutely required for the maintenance of highly proliferative stem and progenitor cell compartments in the intestine, but also for lineage decisions, differentiation and proper positioning of intestinal epithelial cells (IECs) [1]. In agreement with its potent mitogenic effects, aberrant WNT/β-CATENIN pathway activity is a major oncogenic driver in colorectal tumorigenesis [1],[4]. In fact, more than 90 % of human colorectal cancers harbor mutations leading to aberrant activation of WNT/β-CATENIN signaling [5]. Animal studies further indicated that WNT/β-CATENIN pathway activity not only suffices for tumor initiation but might also be essential for tumor maintenance [6], altogether providing strong incentives to interfere with WNT/β-CATENIN signaling for therapeutic purposes [4].

In the healthy intestine, WNT/β-CATENIN pathway-induced transcriptional responses are predominantly if not exclusively mediated by TCF7L2, which is the sole TCF/LEF family member capable to maintain stem and progenitor cell proliferation in vivo [7]–[9], and to sustain viability of small intestinal and colonic organoids in vitro [8]–[11]. Based on the close and essential partnership between β-CATENIN and TCF7L2 in healthy IECs, one would assume that TCF7L2 is indispensable to transmit oncogenic WNT/β-CATENIN pathway activity. However, various findings led to more complex and conflicting views about TCF7L2 and its role in colorectal tumorigenesis. The observation of frequently occurring TCF7L2 loss-of-function mutations in human CRC genomes combined with results from knock-out (KO) studies in a transgenic mouse line and knock-down experiments in certain CRC cell lines pointed to growth inhibitory and tumor suppressor properties of TCF7L2 [5],[12]–[15]. This indicated that CRC cells unlike healthy IECs may not depend on TCF7L2 for survival. Indeed, acquired TCF7L2-independence could experimentally be confirmed by CRISPR/Cas9-mediated inactivation of TCF7L2 in CRC cell lines. However, no support for antiproliferative or tumor suppressive activities of TCF7L2 was found [8],[10]. Rather, in these models TCF7L2-deficiency was accompanied by highly cell-line-specific phenotypic alterations ranging from no discernible effects in some cases to reduced proliferation and enhanced migration and invasion in others [8],[10], the latter being reminiscent of the physiological functions of TCF7L2 in healthy IECs. It thus appears that certain aspects of the function of TCF7L2 in healthy IECs are conserved in CRC cells, but the strict addiction to β-CATENIN/TCF7L2-driven transcription might be lost.

To explain the perplexing TCF7L2-independence of CRC cells it was suggested that other TCF/LEF family members might compensate the loss of TCF7L2 but this has not been proven so far [8],[10],[16]. As an alternative, CRC cell viability might depend on gene expression programs that are driven not by β-CATENIN and TCF/LEF proteins but instead by β-CATENIN and other transcription factors [17]. Further, also complete independence of CRC cells from WNT/β-CATENIN pathway activity was reported [18],[19] which would also explain why a subset of human colorectal tumors and CRC cell lines can tolerate the loss of TCF7L2.

To shed more light on the importance of TCF7L2 and its role in colorectal cancer, we initially aimed to test the idea that other TCF/LEF family members rescue TCF7L2-deficiency. However, the ability to create viable CRC cell lines that do not express any TCF/LEF proteins, demonstrates that TCF/LEF activity is not strictly required for CRC cell survival. This prompted us to extend our analyses to β-CATENIN which likewise turned out to be non-essential in four out of eight CRC cell lines examined. Despite being expendable, β-CATENIN and TCF7L2 still promoted CRC cell proliferation, most likely a vestige of their function in intestinal stem and progenitor cells. Although expression of cognate WNT/β-CATENIN target genes is impaired in β-CATENIN- and TCF/LEF-deficient cells, comprehensive and unbiased analyses of β-CATENIN- and TCF7L2-dependent transcriptomes revealed highly cell-line-specific transcriptional effects and extensive uncoupling of β-CATENIN and TCF7L2 transcriptional activities. Apparently, unlike in the healthy intestinal epithelium, the two factors act largely independently of each other in the context of CRC cells. The pronounced context-dependent and highly distinct, yet non-essential functions of β-CATENIN and TCF/LEF proteins challenge their broad utility as drug targets for CRC therapy.

## Results

### CRC cells are viable in the complete absence of TCF/LEF expression

Previously it was proposed that lethality, which results from TCF7L2-deficiency in healthy IECs, might be prevented in CRC cells due to expression of other TCF/LEF family members that could substitute for TCF7L2 [8],[10]. To test this idea, we attempted to additionally inactivate the *TCF7* and *TCF7L1* genes in HT29 and HCT116 *TCF7L2* single knock-out (SKO) cells (Fig. 1). *TCF7* and *TCF7L2* are the only TCF/LEF family members expressed in HT29 cells, while HCT116 cells express *TCF7, TCF7L1,* and *TCF7L2. LEF1* is not expressed in HT29 and HCT116 cells. (Fig. 1b-e). Therefore, if viable, HT29 *TCF7/TCF7L2* double knock-out (DKO) and HCT116 *TCF7/TCF7L1/TCF7L2* triple knock-out (TKO) cells would not express any TCF/LEF proteins. Strikingly, by using the CRISPR/Cas9 system and a frame-shift inducing exon-deletion strategy we succeeded in generating multiple clonally-derived HT29 DKO and HCT116 TKO cell lines. Compared to their wild type (WT) counterparts, these cells showed much reduced *TCF7, TCF7L1,* and *TCF7L2* transcript levels, and, importantly, had no detectable TCF/LEF protein expression (Fig. 1b-e; Suppl. Tables S1, S2). From this we conclude that *TCF7* and *TCF7L1* do not compensate the loss of *TCF7L2* to maintain CRC cell viability. Rather, it appears that TCF/LEF expression is entirely dispensable for the survival of HT29 and HCT116 cells.

**Figure 1.**
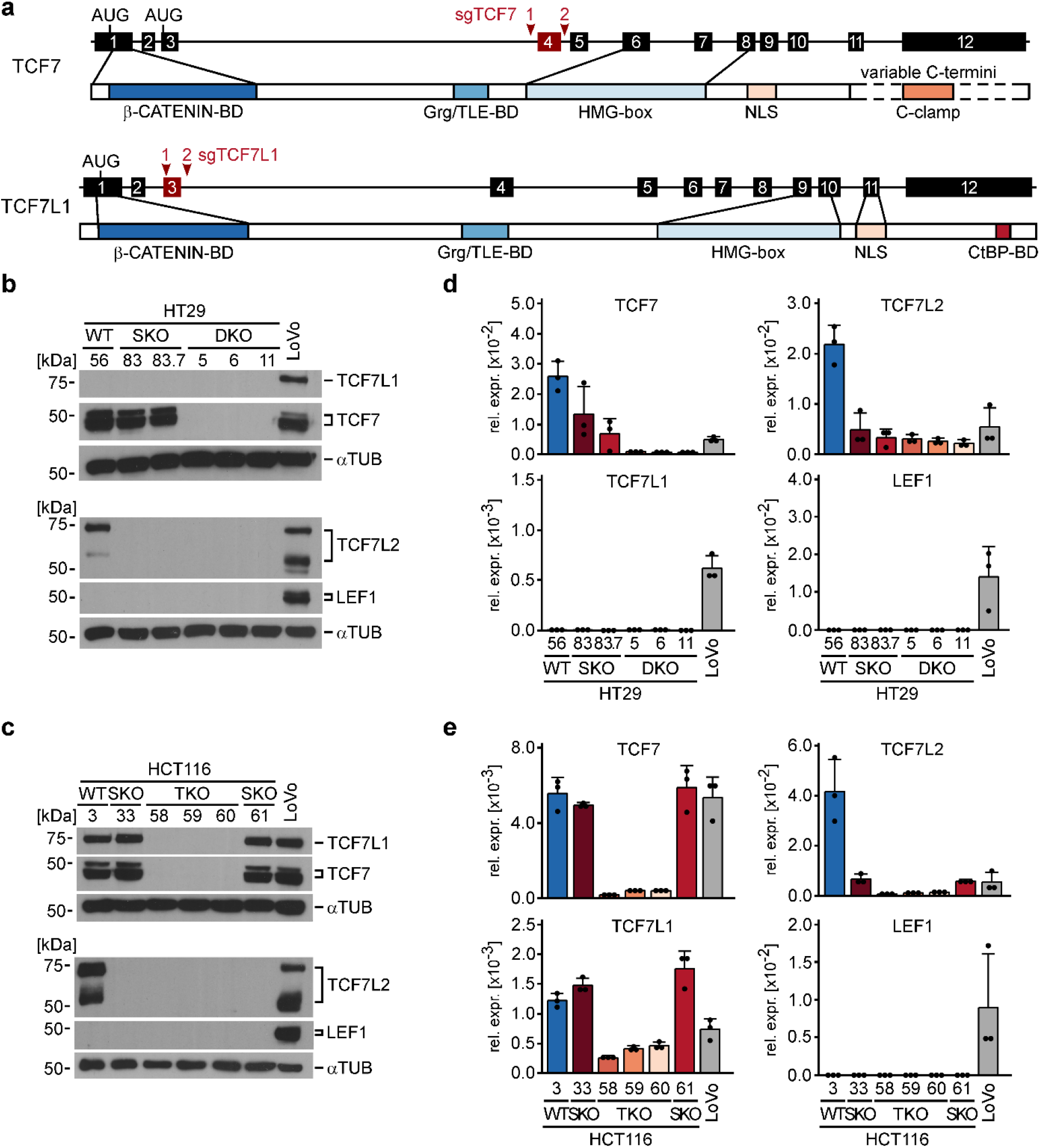
HT29 and HCT116 CRC cell lines are viable in the complete absence of TCF/LEF transcription factors. **a** Schematic representation of the *TCF7* and *TCF7L1* genes and their protein products. Exons are depicted as numbered boxes. Exons chosen for CRISPR/Cas9-induced deletion are shown in red. The locations of sgRNA target sites are indicated by red arrowheads. The position of start codons (AUG) and functionally important domains of the TCF7 and TCF7L1 proteins are marked and connected to the exon(s) by which they are encoded. HMG-box: DNA-binding domain; NLS: nuclear localization sequence; Grg: Groucho-related gene; TLE: Transducin-like enhancer of split; CtBP: Carboxy-terminal binding protein; BD: binding domain. **b**, **c** Western blot analyses to detect TCF/LEF expression in whole cell lysates from HT29 (**b**) and HCT116 cells (**c**) with the genotypes indicated. WT: wild type; SKO: single knock-out, biallelic inactivation of *TCF7L2*; DKO: double knock-out, HT29 cells with biallelic inactivation of *TCF7* and *TCF7L2.* TKO: triple knock-out, HCT116 cells with biallelic inactivation of *TCF7, TCF7L1, and TCF7L2.* LoVo cells were included as positive control for TCF/LEF detection. α-TUBULIN (αTUB) was used to demonstrate equal loading. Molecular weights are given in kDa. Representative results from one of three independent biological replicates are shown. Full size/uncropped versions of the Western blot images are shown in Suppl. Fig. S9. **d**, **e** The bar plots summarize and compare RNA expression levels of TCF/LEF family members determined by qRT-PCR in HT29 (**d**) and HCT116 cells (**e**) with the genotypes indicated. LoVo cells were included as positive control. TCF/LEF transcript levels were normalized to those of *GAPDH*, and are displayed as relative expression (rel. expr.). Each dot represents an individual measurement. Error bars indicate SD (n=3).

### Redundant and non-redundant functions of TCF/LEF family members in HT29 and HCT116 CRC cell lines

Even though TCF7 and TCF7L1 apparently do not play a role in acquired TCF7L2-independence of CRC cells, we nonetheless asked whether their additional absence had an impact on certain phenotypic changes observed in *TCF7L2* SKO cells [10]. First, we examined cellular morphology. We found that HT29 SKO and HT29 DKO cells both exhibited a loss of the densely clustered and cobblestone-like growth pattern displayed by a representative HT29 WT cell clone. However, HT29 SKO and DKO cells did not noticeably differ from each other (Fig. 2a). In contrast, the elongated, spindle-like shape of HCT116 WT cells clearly changed only in the complete absence of TCF/LEF expression. Compared to HCT116 WT and SKO cells, HCT116 TKO cells grew more disperse and acquired roundish cell shapes, similar to HT29 SKO and DKO cells (Fig. 2a). Apparently, TCF7L2 has a unique impact on HT29 cell morphology whereas in HCT116 cells TCF7, TCF7L1, and TCF7L2 may function interchangeably or in a combinatorial fashion to determine cellular appearance. Likewise, analyses of WNT/β-CATENIN target gene expression argues for cell-type-specific overlapping as well as distinct functions of TCF/LEF family members. Thus, in HT29 cells, the sole absence of TCF7L2 already reduced expression of *AXIN2, RNF43, MYC,* and *TERT.* Except for *AXIN2,* their expression did not further decrease in TCF7/TCF7L2 double-deficient cells (Fig. 2b). In HCT116 cells, however, single loss of TCF7L2 impaired expression of *RNF43* and *MYC*, but not of *AXIN2* and *TERT,* the expression of which was diminished only in HCT116 TKO cells (Fig. 2b). Thus, *RNF43* and *MYC* appear to be under unique control by TCF7L2 while *AXIN2* and to some extent *TERT* are additionally regulated by TCF7 and/or TCF7L1. Finally, we analyzed whether defects in cell proliferation, which we had observed in TCF7L2-deficient cells [10], were affected by combined loss of TCF/LEF family members. Yet, cell counting and flow cytometry experiments revealed that impaired population dynamics and G1/S phase progression shown by HT29 and HCT116 SKO cells were not further aggravated in HT29 DKO and HCT116 TKO cells (Fig. 2c, Suppl. Fig. S1). In aggregate, these observations argue for context-dependent, combinatorial and partially redundant functions of TCF/LEF family members, for example in the regulation of cell shape and expression of some target genes, but also for non-redundant roles of TCF7L2 as in cell cycle control which is in line with its unique requirement in IEC proliferation.

**Figure 2.**
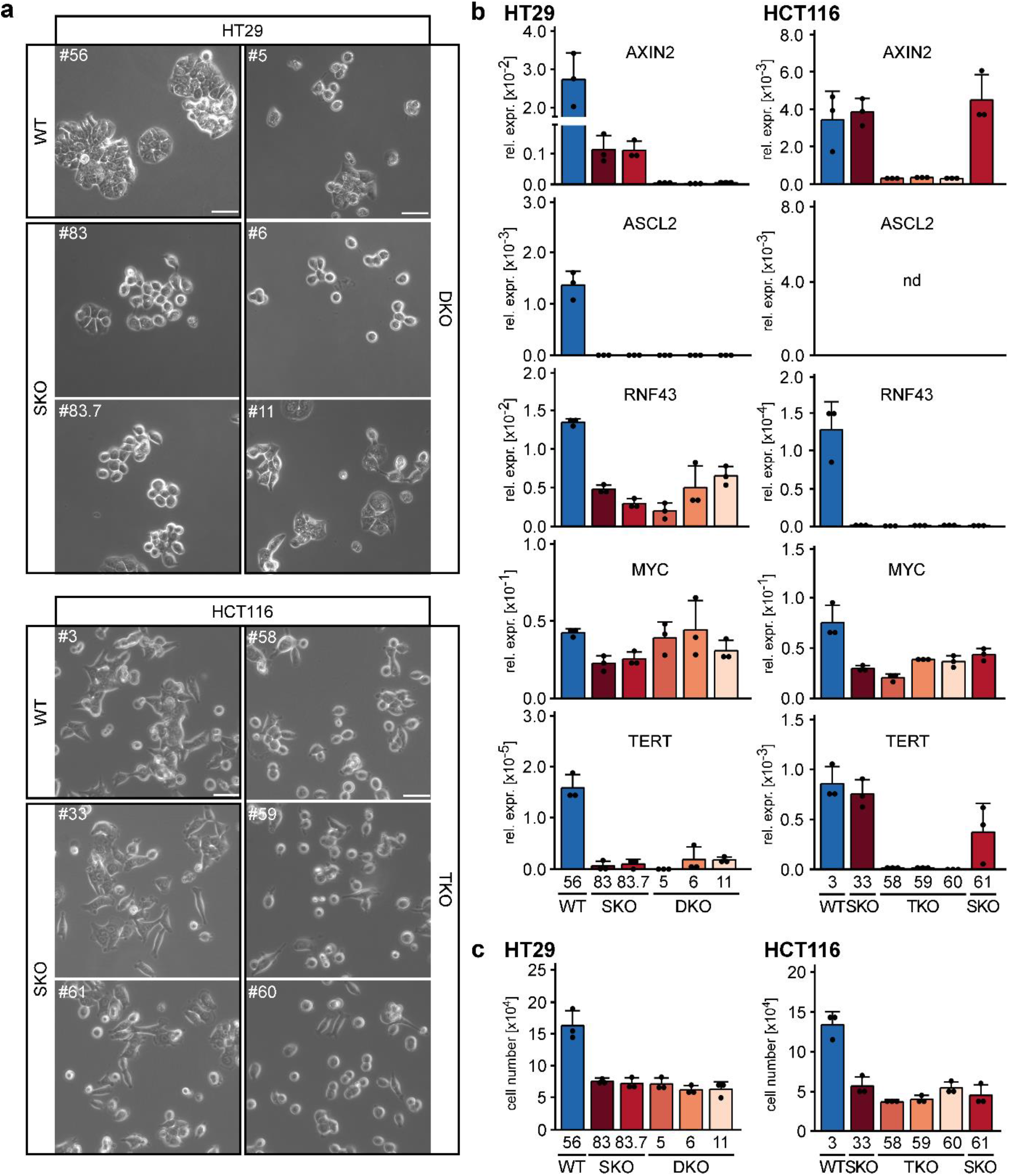
Redundant and non-redundant functions of TCF/LEF family members in HT29 and HCT116 CRC cell lines. **a** Representative micrographs from one of three independent biological replicates showing the morphology of HT29 and HCT116 WT, SKO, DKO, and TKO cells 24 h after seeding the same starting numbers of cells. The scale bars represent 50 μm. **b** Expression of the WNT/β-CATENIN target genes *AXIN2, ASCL2, RNF43, MYC,* and *TERT* was analyzed by qRT-PCR in HT29 and HCT116 cell clones with the genotypes indicated. Transcript levels were normalized to those of *GAPDH,* and are displayed as relative expression (rel. expr.). The bar plots summarize the expression data; error bars indicate SD (n=3). Each dot represents an individual measurement. nd: not detectable. **c** To compare population dynamics of WT and TCF/LEF-deficient cells, 5×10^4^ HT29 and HCT116 cells with the genotypes indicated were seeded, incubated for 72 h and counted. The resulting cell counts are displayed as bar plots with each dot representing an individual measurement. Error bars indicate SD (n=3).

### CRC cells can survive the loss of β-CATENIN

While β-CATENIN can also interact with other DNA-binding proteins to mediate transcriptional responses [17], TCF/LEF family members are considered to be the major if not exclusive factors mediating WNT/β-CATENIN pathway-induced transcriptional responses [20]. The finding that HT29 and HCT116 cells survive in the absence of any TCF/LEF expression therefore raised the question whether CRC cells also exhibited independence from β-CATENIN. To address this, we tested whether viable CRC cell line derivatives could be obtained when targeting the *CTNNB1* gene (coding for β-CATENIN) using CRISPR/Cas9 technology. The knock-out attempt was made in HT29, HCT116, LoVo, LS174T, LS411, SW403, SW480, and RKO cells. Except for RKO cells, all other cell lines tested have inactivating mutations in *APC* (HT29, LoVo, LS411, SW403, SW480) or gain-of-function mutations in *CTNNB1* (HCT116, LS174T), which lead to growth factor-independent WNT/β-CATENIN pathway activation and transcriptional activity. Surprisingly, using the exon deletion strategy outlined in Fig. 3a, we were able to generate several clonal *CTNNB1*^-/-^ cell lines from HCT116, HT29, SW480, and RKO cells that showed strongly reduced *CTNNB1* RNA levels and no β-CATENIN protein expression (Suppl. Fig. S2a-d; Fig. 3b; Suppl. Tables S1, S2). No clones with biallelic inactivation of *CTNNB1* were retrieved from LoVo, LS174T, LS411, and SW403 cells although it is unclear whether this is because of technical issues or due to essential functions of β-CATENIN in these cells. Similar to TCF/LEF-deficiency, also loss of β-CATENIN had cell-line-specific effects on cell morphology (Fig. 3c). *CTNNB1*^-/-^ HT29 and SW480 cells appeared larger, flattened, and grew in less compact cell clusters compared to *CTNNB1*^+/+^ cells. Yet, *CTNNB1*^-/-^ HT29 cells showed clonal variability with respect to the severity of morphological changes. In contrast, *CTNNB1*^-/-^ HCT116 and RKO cells showed no obvious changes in cell shape. On the other hand, as further difference among *CTNNB1*^-/-^ CRC cells, we noticed that upon prolonged cultivation, *CTNNB1*^-/-^ SW480 cell clones became senescent, indicated by proliferation arrest, vastly increased cell size, and pronounced β-GALACTOSIDASE activity (Suppl. Fig. S3). However, senescent *CTNNB1*^-/-^ SW480 cells seemingly remained alive for several weeks (Suppl. Fig. S3). Importantly, acquisition of a senescent phenotype was not observed with *CTNNB1*^-/-^ HT29, HCT116, and RKO cells, and did not correlate with WT and mutant states of *TP53* [21]. Altogether, we conclude that there is no obligatory requirement for β-CATENIN for viability of CRC cell lines.

**Figure 3.**
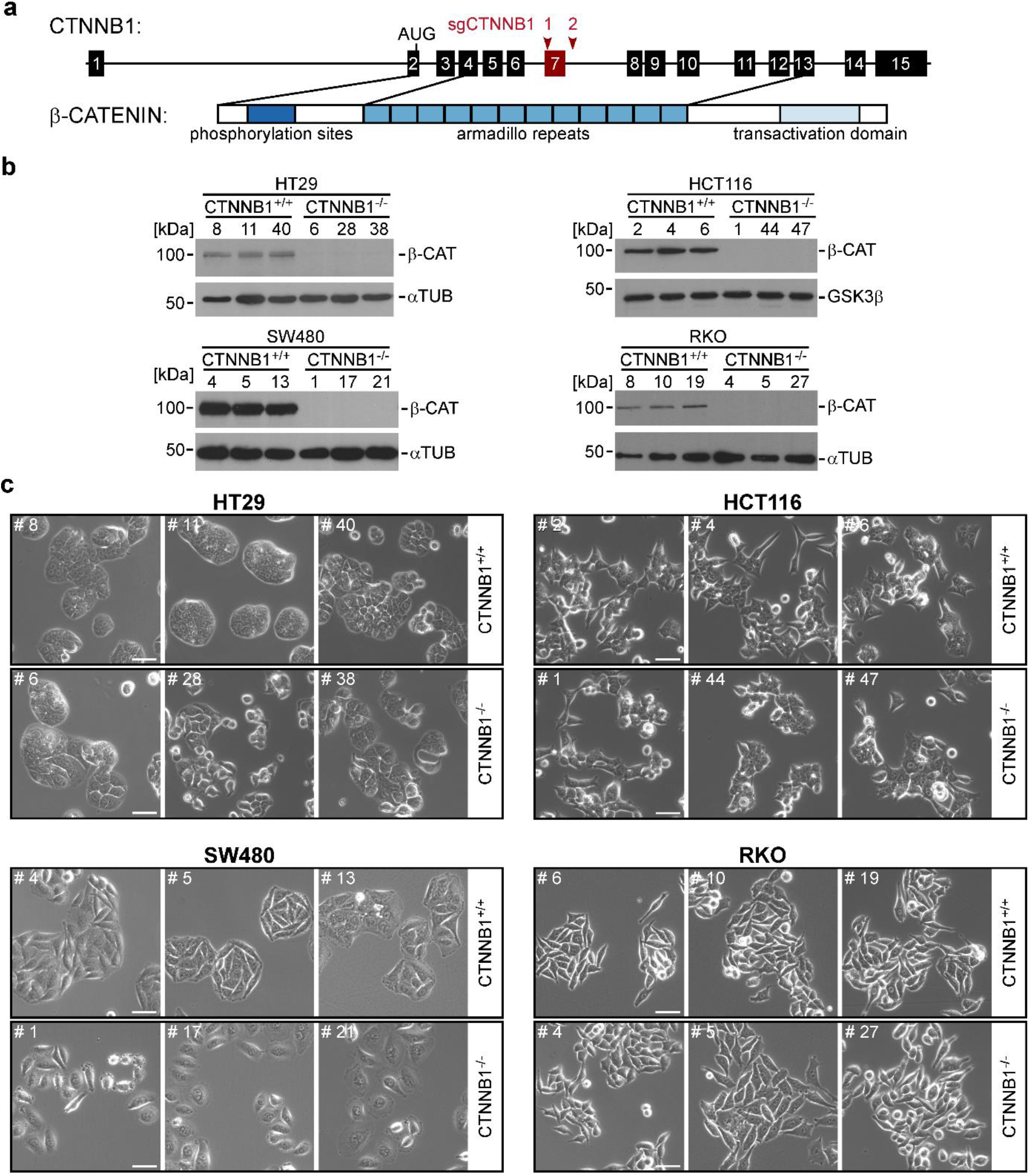
Colorectal cancer cell lines are viable without β-CATENIN. **a** Schematic representation of the *CTNNB1* gene and its protein product β-CATENIN. Exons are depicted as numbered boxes. Exon 7 (red box) was chosen for CRISPR/Cas9-induced deletion using two sgRNAs targeting positions marked by red arrowheads. The location of the start codon (AUG) in exon 2 and functionally important domains of β-CATENIN are indicated and connected to the exons by which they are encoded. **b** Western blot analyses to detect β-CATENIN (β-CAT) expression. Cell lysates were generated from cells with biallelic WT and mutant *CTNNB1* genes. α-TUBULIN (αTUB) or GSK3β were used as loading controls. Molecular weights are given in kDa. Representative results from one of three independent biological replicates are shown. Full size/uncropped versions of the Western blot images are shown in Suppl. Fig. S9. **c** Representative micrographs from one of three independent biological replicates showing HCT116, HT29, SW480 and RKO cell clones with biallelic WT and mutant *CTNNB1* genes. Images were taken 24 h after seeding the same starting numbers of cells. The scale bars represent 50 μm.

### β-CATENIN-deficiency abolishes transcriptional activity of WNT/β-CATENIN signaling

To assess the impact of β-CATENIN-deficiency on transcriptional activity of the WNT/β-CATENIN pathway, we first performed reporter gene assays using the pSuper8xTOPFlash and pSuper8xFOPFlash pair of luciferase expression vectors with WT and mutant TCF/LEF binding sites in their promoter regions, respectively [22]. The two reporter constructs were transiently transfected into HT29, HCT116, pre-senescent SW480, and RKO cells. Comparison of the luciferase activities produced by the WT and mutant reporter plasmids confirmed the presence of β-CATENIN and TCF/LEF-mediated transcriptional activity in *CTNNB1^+/+^* HT29, HCT116, and SW480 cells, which was completely abolished in *CTNNB1*^-/-^ cells (Suppl. Fig. S4). In agreement with the WT state of their *APC* and *CTNNB1* genes and the absence of other mutations that could activate the WNT pathway, RKO cells showed no intrinsic β-CATENIN transcriptional activity. Still, upon treatment with the GSK3β inhibitor CT99021, luciferase expression from the WT reporter increased in *CTNNB1*^+/+^ RKO cells. As expected, the reporter response was completely blocked in *CTNNB1*^-/-^ RKO cells (Suppl. Fig. S4).

Next, we investigated the impact of β-CATENIN-deficiency on the expression of the cellular target genes *AXIN2, ASCL2, RNF43, MYC,* and *TERT.* In cell lines with an active WNT/β-CATENIN pathway (HT29, HCT116, SW480) and provided that the gene under investigation was expressed, we observed that eliminating β-CATENIN was paralleled by reduced expression of *AXIN2, ASCL2,* and *TERT,* albeit the reduction in *TERT* expression did not reach statistical significance in HT29 cells (Fig. 4). Likewise, *RNF43* expression was lower in *CTNNB1*^-/-^ HCT116 and SW480 cells. Surprisingly, however, *RNF43* expression was not affected by β-CATENIN-deficiency in HT29 cells even though it had proven to be under control by TCF7L2. Similarly, and also in contrast to the response to TCF/LEF depletion, *MYC* transcript levels showed only a small, and statistically non-significant decline in β-CATENIN-deficient HCT116 and SW480 cells, and even went up in *CTNNB1*^-/-^ HT29 cells (Fig. 4a). The latter was also seen in RKO cells (Fig. 4d). On the other hand, *AXIN2* and *TERT* expression was not affected when *CTNNB1* was inactivated in RKO cells, which is in line with the resting state of the WNT/β-CATENIN pathway in these cells. Taken together, although the reporter gene assays had demonstrated that β-CATENIN and TCF/LEF-mediated transcriptional activity was effectively and uniformly abolished in β-CATENIN-deficient cells, endogenous, WNT/β-CATENIN target genes showed more diverse and cell-line specific differences with respect to their overall expression and response to ablation of β-CATENIN.

**Figure 4.**
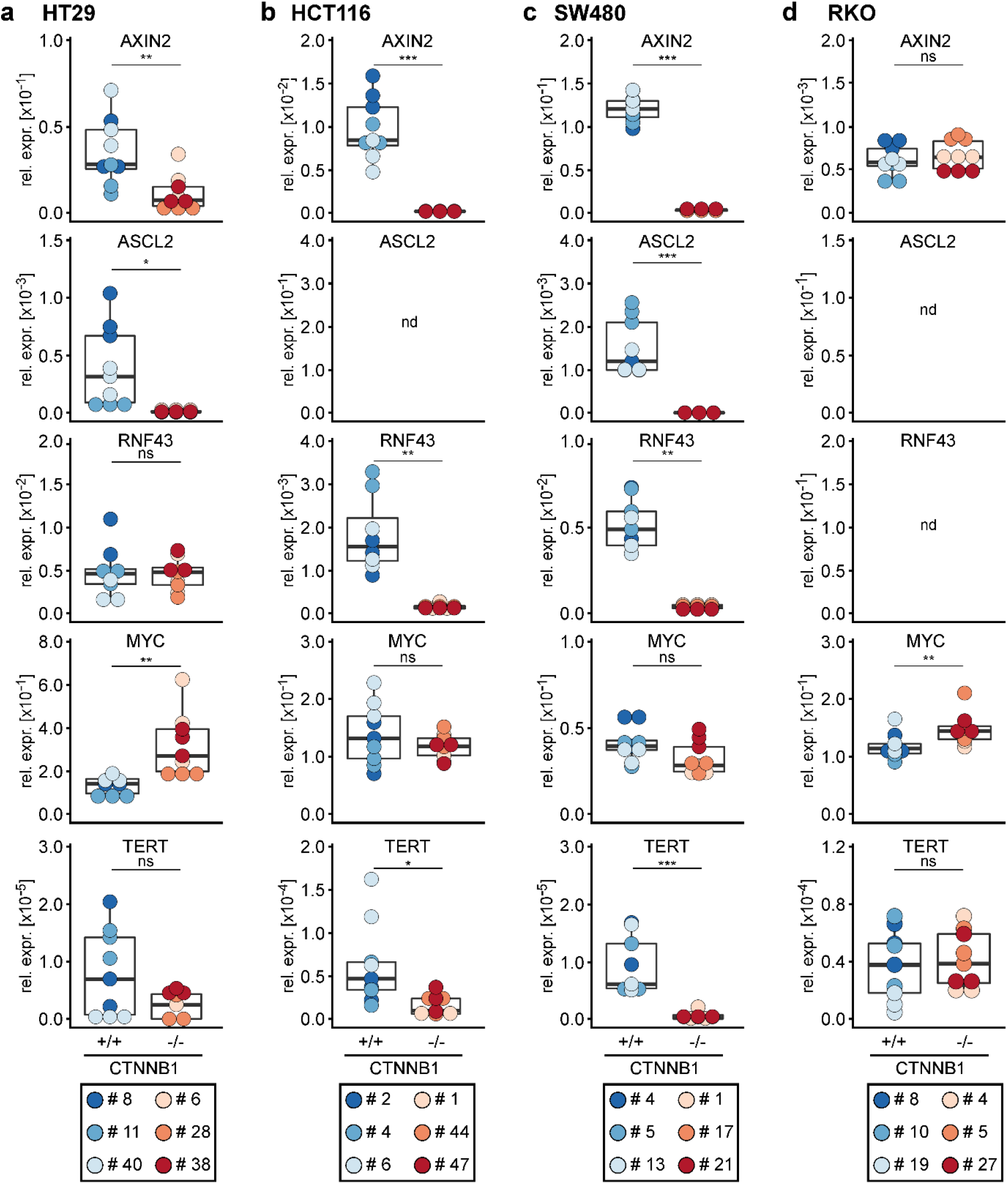
WNT/β-CATENIN target gene expression is reduced in β-CATENIN-deficient CRC cells. **a-d** Expression of the WNT/β-CATENIN target genes *AXIN2, ASCL2, RNF43, MYC,* and *TERT* was analyzed by qRT-PCR in HT29 (**a**), HCT116 (**b**), SW480 (**c**) and RKO (**d**) cell clones with biallelic WT and mutant *CTNNB1* genes. Transcript levels were normalized to *GAPDH,* and are displayed as relative expression (rel. expr.). The box plots display aggregated expression data from *CTNNB1*^+/+^ and *CTNNB1*^-/-^ cell clones. Each dot represents a separate measurement with dot color identifying individual clones. For statistical analysis, LMM was performed (n=3). nd: not detectable; ns: not significant.

### β-CATENIN-deficiency affects proliferation and cell cycle progression in some CRC cells

WNT/β-CATENIN signaling is a potent mitogenic pathway in intestinal stem and progenitor cells as well as intestinal tumor cells, and blocking the pathway by inactivating *TCF7L2* impairs CRC cell cycle progression (see above and [10]). Therefore, we were curious to see whether β-CATENIN-deficiency likewise affected proliferation of CRC cells. Indeed, by seeding defined numbers of cells and counting their progeny 72 h later, we found that relative to their WT counterparts β-CATENIN-deficient HT29, HCT116, and SW480 cells displayed reduced population dynamics (Fig. 5) which was not due to increased apoptosis (Suppl. Fig. S5). Rather, analyses of cellular DNA content by flow cytometry suggested a delay in G1/S phase transition in *CTNNB1*^-/-^ HT29 and HCT116 cells (Fig. 5b), which had also been seen in TCF7L2-deficient cells (see above and [10]). In contrast, the clearly retarded expansion of SW480 cell populations could not be linked to a defined block in cell cycle progression. Besides, and as expected, proliferation of RKO cells with an inactive WNT/β-CATENIN pathway did not change in the absence of β-CATENIN. In sum, it appears that the mitogenic effects of WNT/β-CATENIN pathway activity found in healthy IECs are partly retained in HT29, HCT116, and SW480 CRC cells, but the strict dependence on β-CATENIN and TCF7L2 for proliferation and survival appears to be alleviated in the cancer cells.

**Figure 5.**
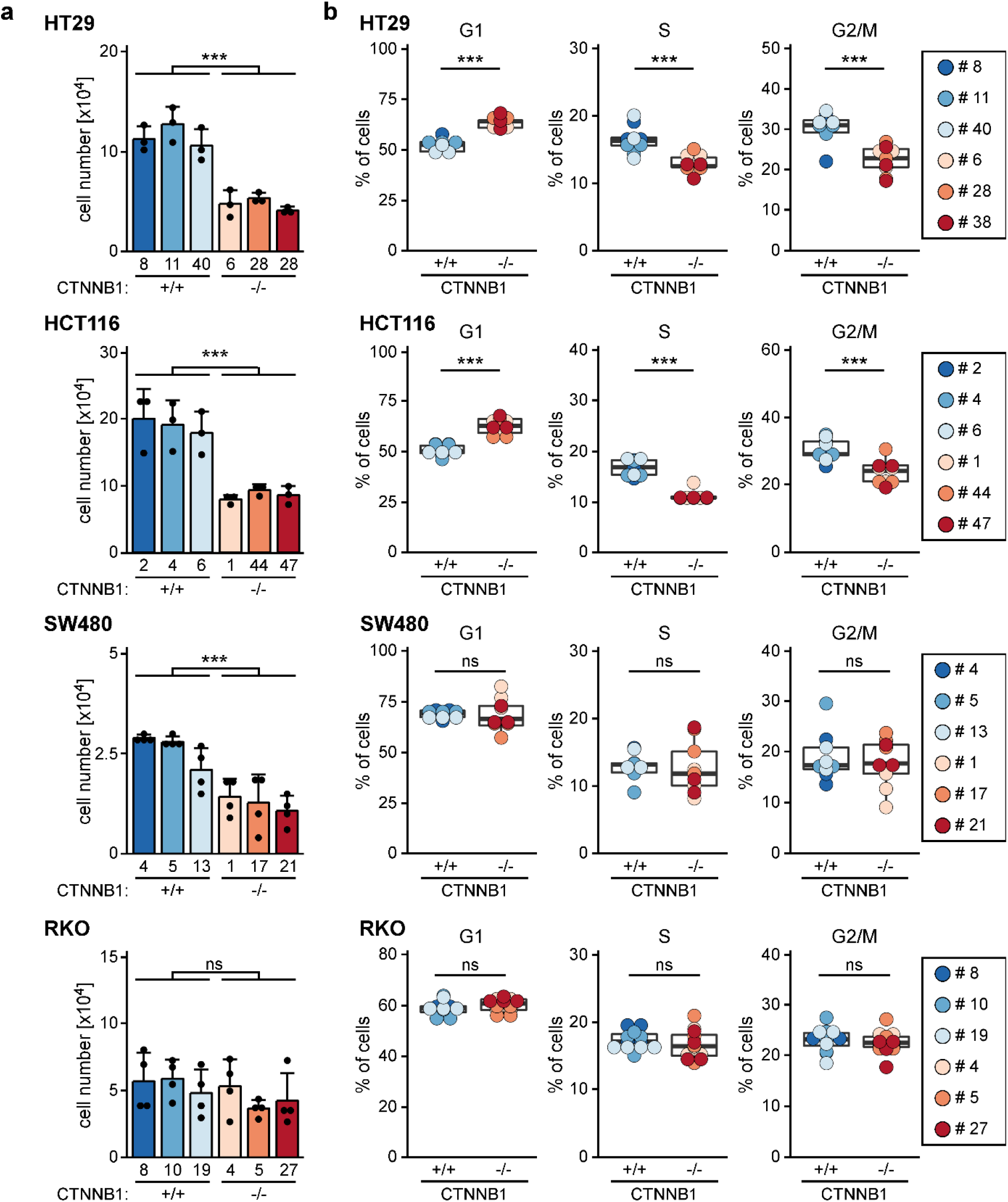
Cell-type-specific impairment of population dynamics and cell cycle progression upon loss of β-CATENIN. **a** To compare population dynamics of WT and β-CATENIN-deficient cells, 1×10^4^ each of HT29, HCT116, SW480, and RKO cells with the genotypes indicated were seeded, incubated for 72 h, and counted. The resulting cell counts are displayed as bar plots whereby each dot represents an individual measurement. For statistical analysis LMM was performed, the error bars indicate SD (n=3). ns: not significant. **b** To analyze differences in cell cycle distribution, HT29, HCT116, SW480, and RKO cells with the genotypes indicated were stained with propidium iodide and analyzed by flow cytometry. The proportions of cells in different cell cycle phases are depicted by box plots. Each dot represents a separate measurement with dot color identifying individual clones. For statistical analysis LMM was performed (n=3). ns: not significant.

### PLAKOGLOBIN may preserve E-CADHERIN membrane localization but cannot sustain WNT target gene expression in β-CATENIN-deficient cells

Aside from its function as transcriptional coactivator in WNT signaling, β-CATENIN also plays a critical role in cell-cell adhesion where it affects intracellular trafficking and stability of CADHERIN transmembrane proteins and is essential for functionality of adherens junctions [23]. HT29 cells have an epithelial morphology and express E-CADHERIN. The observed clonal variability notwithstanding, we therefore were surprised that the loss of β-CATENIN did not more severely alter the growth pattern of HT29 cells. Accordingly, we tested whether absence of β-CATENIN affected the localization of E-CADHERIN by immunofluorescence staining. However, E-CADHERIN membrane staining was maintained in *CTNNB1*^-/-^ HT29 cells (Suppl. Fig. S6). Previous work suggested that the closely related PLAKOGLOBIN (also known as γ-CATENIN) in some cases may substitute for β-CATENIN to uphold cell-cell adhesion [24]–[26]. Indeed, HT29 cells express PLAKOGLOBIN which appears to colocalize with E-CADHERIN at cell-cell interfaces in *CTNNB1*^+/+^ as well as *CTNNB1*^-/-^ cells (Suppl. Figs. S7, S8). Co-immunoprecipitation experiments confirmed that E-CADHERIN forms complexes not only with β-CATENIN but also with PLAKOGLOBIN in HT29 cells (Suppl. Fig. S8). This interaction was not affected by the presence or absence of β-CATENIN. Thus, PLAKOGLOBIN may maintain E-CADHERIN membrane localization and possibly adhesive function when β-CATENIN expression is abrogated. Yet, in view of the results of our reporter gene studies, PLAKOGLOBIN seemingly cannot take over the function of β-CATENIN in WNT target gene activation. This conclusion is in agreement with earlier reports of differential gene regulatory capacities of PLAKOGLOBIN and β-CATENIN [27]–[29].

### Transcriptome analyses confirm highly cell-line-specific effects of β-CATENIN-deficiency

Targeted gene expression analyses had revealed differential, cell-line-specific alterations ensuing from β-CATENIN-deficiency. To capture differences in the transcriptomes of *CTNNB1*^+/+^ and *CTNNB1*^-/-^ cells in an unbiased and comprehensive way, we performed two independent rounds of RNA-seq experiments for two *CTNNB1*^+/+^ and *CTNNB1*^-/-^ clones each derived from the HT29, HCT116, and SW480 cell lines. Principal component analyses (PCA) showed a high degree of similarity between the two replicates for each cell clone (Fig. 6a). Further, for HCT116 and SW480 cells, *CTNNB1*^+/+^ and *CTNNB1*^-/-^ clones were well separated along principal component 1 which covers the largest part of the variance and likely reflects WT and mutant states of *CTNNB1.* Consistent with the observed clonal variability, PCA indicated considerable differences among HT29 cell clones which cannot be explained solely by their genotypes. Irrespective of this, by comparing gene expression profiles of *CTNNB1*^+/+^ and *CTNNB1*^-/-^ cell clones and by using an absolute log2(fold change)>1 and an adjusted p-value<0.05 as thresholds, we identified 868 differentially expressed genes (DEGs) in HT29 cells, 779 DEGs in HCT116 cells, and 3,581 DEGs in SW480 cells (Fig. 6b,c; Suppl. Tables S3-S5). In each cellular background, the majority of DEGs were upregulated in *CTNNB1*^-/-^ cell clones relative to the *CTNNB1*^+/+^ control cells (Fig. 6c; Suppl. Tables S3-S5). Gene set enrichment analyses based on these DEGs and the collection “Biological process” pointed to various gene ontology (GO) terms related to RNA processing, morphogenesis, organogenesis and β-CATENIN-TCF complex assembly as being most negatively enriched in β-CATENIN-deficient cells (Suppl. Tables S6-S8). Nonetheless, comparison of DEGs across cell lines corroborated that β-CATENIN-deficiency produced highly cell-line specific changes in gene expression in HT29, HCT116, and SW480 cells (Fig. 6b). This became even more apparent when taking into account the directionality of regulation (Fig. 6c). Altogether, only 51 DEGs were common to HT29, HCT116, and SW480 cells (Fig. 6b, Suppl. Table S9). Notably, aside from *CTNNB1* and *AXIN2,* genes downregulated in all three *CTNNB1*^-/-^ cell types included *NR2F1* (COUP-TFII), *ADAM19,* and *BMP4* which are also known WNT/β-CATENIN target genes [30]–[34]. Nevertheless, the majority of the 51 shared DEGs were upregulated in *CTNNB1*^-/-^ cells and quite often were regulated in opposite directions in the three cell lines. In their sum, the results of the transcriptome analyses therefore revealed an extensive diversification of β-CATENIN-dependent gene expression in CRC cell lines.

**Figure 6.**
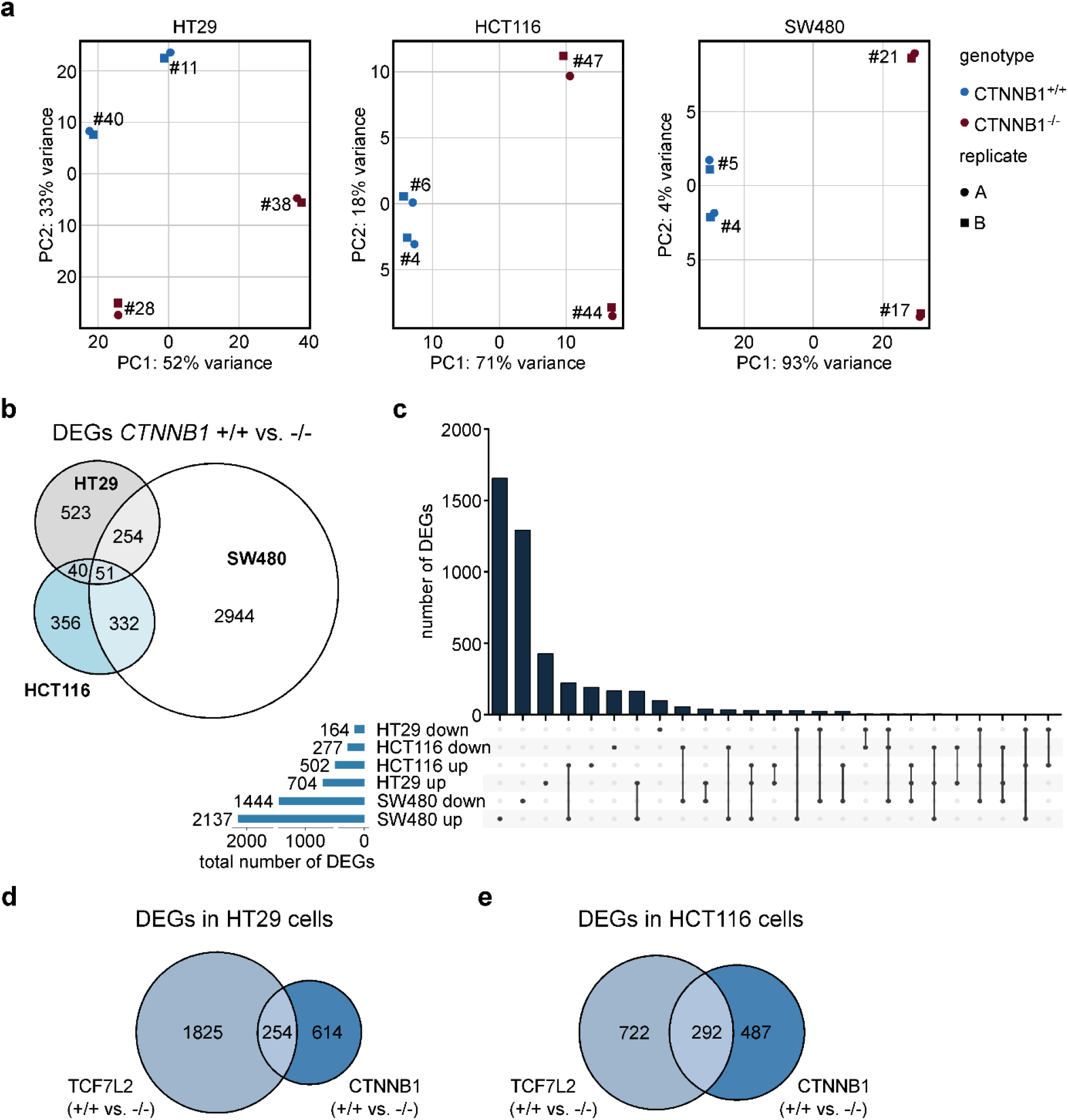
Loss of *CTNNB1* results in highly cell-type-specific gene expression changes in CRC cell lines. **a** Principal component analysis (PCA) of transcriptome data derived from HT29, HCT116, and SW480 cell clones with biallelic WT and mutant *CTNNB1* genes. For each cell line and genotype, two different cell clones were analyzed by RNA-seq. Two independent biological replicates were performed (n=2). **b** Venn diagram showing the numbers of differentially expressed genes (DEGs) specific and common to HCT116, HT29, and SW480 cells with biallelic WT and mutant *CTNNB1* genes. **c** Upset plot depicting the numbers of DEGs specific and common to HCT116, HT29, and SW480 cells with biallelic WT and mutant *CTNNB1* genes taking into account the direction of their regulation. In addition, for each cellular background the total numbers of up- and downregulated DEGs are given. **d**, **e** Comparison of gene expression changes following inactivation of *TCF7L2* and *CTNNB1* in HT29 (**d**) and HCT116 cells (**e**). The Venn diagrams show the numbers of DEGs specific and common to cells with biallelic WT and mutant *TCF7L2* or *CTNNB1* genes. **b-e** As thresholds to call DEGs, absolute values of a log2(fold change)>1 and an adjusted p-value<0.05 were used.

The pronounced cell-line-specificity of transcriptomic changes caused by β-CATENIN-deficiency were reminiscent of similar observations made with TCF7L2-deficient cell lines [10]. It was therefore of interest to examine whether β-CATENIN and TCF7L2 regulated the same or divergent sets of genes in a given cellular background. Remarkably, in both HT29 and HCT116 cells, TCF7L2 appeared to control more genes than β-CATENIN (Fig. 6d,e). Furthermore, only 29 % and 37 % of the genes regulated by β-CATENIN in HT29 and HCT116 cells, respectively, were also under control of TCF7L2 demonstrating that β-CATENIN and TCF7L2 control sizeable fractions of their target genes independently from each other. Thus, HT29 and HCT116 CRC cells do not feature the close partnership between β-CATENIN and TCF7L2 in gene regulation that is a hallmark of healthy IECs.

## Discussion

Proliferation and survival of healthy intestinal stem and progenitor cells critically depend on the activity of the WNT/β-CATENIN pathway and its essential effectors β-CATENIN and TCF7L2 [7]–[9],[35],[36]. Despite the mutual dependence of β-CATENIN and TCF7L2 in healthy IECs and for tumor initiation, high proportions of colorectal cancer genomes carry inactivating mutations in the *TCF7L2* gene [5],[13],[15],[37]. Further, tumor suppressor and growth inhibitory functions were ascribed to *TCF7L2* [12],[14]. These observations suggested that *TCF7L2* activity may not be mandatory for survival of colorectal tumor cells, which was experimentally confirmed in several distinct CRC cell lines [8],[10],[16]. To explain the differential requirements for *TCF7L2* in healthy IECs and in CRC cells it was hypothesized that redundancy among TCF/LEF family members may allow for the loss of TCF7L2 in a cancer context [8],[10]. Here, we tested this idea and report that survival of at least some CRC cell lines does not require any TCF/LEF expression, and that several randomly chosen CRC cell lines even tolerate inactivation of β-CATENIN. Thus, it appears that the vital dependence on β-CATENIN and TCF/LEF transcriptional activity that characterizes normal gut tissue, can be lost during colorectal carcinogenesis and that this may not be an exceptional and rare event. Nonetheless, in the absence of β-CATENIN or TCF/LEF proteins, CRC cell lines suffer from mitotic defects, arguing that they retain to varying degrees some vestiges of the function of β-CATENIN and TCF7L2 in normal tissue.

The idea that CRC cells bear the loss of TCF7L2 because other TCF/LEF family members substitute for it, was sparked by reports of functional interchangeability among TCF/LEF family members [38],[39] and observations that TCF/LEF expression patterns in CRC cell lines and colorectal tumors differ from those in normal IECs [8],[10],[40],[41]. This includes for instance aberrant upregulation of LEF1 [10],[40],[41]. However, the results of our current study clearly refute this theory and actually demonstrate that none of the TCF/LEF proteins are essential for CRC viability. In retrospect, the finding that it is not expression of other TCF/LEF family members which prevents lethality caused by inactivation of TCF7L2, is not too surprising.

TCF7 and TCF7L1 are coexpressed with TCF7L2 in healthy IECs [9],[10],[42], yet fail to complement the TCF7L2 knockout in vivo [7]–[9] and in vitro [8]–[11]. Furthermore, several previous studies showed that TCF7L2 and LEF1 have very different transcription regulatory properties and rather antagonize each other [43]–[45] which argues that also LEF1 is unlikely to take over the function of TCF7L2. As a consequence, we rule out that cancer-specific neoexpression and compensation by other TCF/LEF family members account for expendability of TCF7L2 and suggest that this much more likely reflects far-reaching WNT/β-CATENIN pathway independence of CRC cell lines.

In this context it is important, though, to distinguish between a vital requirement for TCF7L2 or any of the other TCF/LEF family members which seemingly can be lost in CRC cell lines, and their impact on gene expression of CRC cells which is clearly detectable. There is no doubt that the cell-line-specific complement of TCF/LEF proteins does influence gene expression profiles of CRC cells and changes in TCF/LEF expression are accompanied by transcriptomic changes [41],[43],[46]. This appears to entail unique but also redundant and combinatorial as well as antagonistic activities and seemingly occurs in a manner that is highly dependent on cellular background and the target genes examined as shown here and in other studies [8],[10],[16],[41],[43],[46].

Overall, we and others successfully inactivated TCF7L2 in five different CRC cell lines [8],[10],[16]. Further, four out of eight CRC cell lines could cope with β-CATENIN-deficiency. Our lack of success in obtaining *CTNNB1*^-/-^ derivatives from LS174T, LS411, LoVo, and SW403 cells could have technical reasons such as low transfection efficiency or insufficient expression of Cas9 and sgRNAs. However, using experimental procedures analogous to those employed here, we could readily generate *LEF1*^-/-^ LS174T cells [43]. This demonstrates the ability to disable a WNT/β-CATENIN pathway component by CRISPR/Cas9 technology in this cell line, and hints at a fundamental difference between the functional importance of LEF1 and β-CATENIN in the LS174T cell background. Indeed, completely blocking WNT/β-CATENIN signaling by expression of either dominant-negative TCF7 or TCF7L2 and, alternatively, by shRNA-mediated knockdown of β-CATENIN, resulted in growth arrest of LS174T cells [47],[48]. Accordingly, we do not think that all CRC cell lines - and by extrapolation all colorectal cancers - lose dependency on WNT/β-CATENIN pathway activity mediated by TCF/LEF proteins and β-CATENIN.

The CRISPR/Cas9 system was used before to generate viable cells entirely negative for TCF/LEF expression [39],[49] or β-CATENIN [49]. However, these earlier studies employed murine embryonic stem cells and HEK293 human embryonic kidney cells which possess no endogenous WNT/β-CATENIN pathway activity and for which no essential functions of WNT/β-CATENIN signaling are known [39],[49]. In contrast, WNT/β-CATENIN signaling is active in CRC cells and the cells are derived from a tissue that exhibits extraordinary dependence on WNT/β-CATENIN pathway activity and its mediators β-CATENIN and TCF7L2 [1],[7]–[10],[12],[35],[36]. The ability to generate viable CRC cell lines without any TCF/LEF or β-CATENIN expression, therefore, is remarkable, and raises the question as to the underlying cause for the observed expendability of β-CATENIN and TCF7L2. The extreme reliance on mitogenic WNT/β-CATENIN pathway activity displayed by the healthy intestine, may to variable degrees be alleviated by oncogenic MAPK or PI3 kinase signaling in colorectal tumors [50]. Aside from this, it was already shown that CRC cells and oncogenically transformed intestinal organoids can lose WNT/β-CATENIN pathway addiction owing to Hh/GLI signaling [18] or nuclear YAP activity, respectively [19]. Currently, however, it is not known whether all CRC cell lines that allow for inactivation of TCF7L2 or β-CATENIN share as common property high Hh/GLI or YAP activity, or whether other, yet to be identified conditions, can render β-CATENIN and TCF7L2 dispensable.

In healthy IECs, β-CATENIN and TCF7L2 drive common or highly congruent transcriptional programs [20]. In contrast, the comparative transcriptome analyses conducted with CRC cell lines lacking β-CATENIN or TCF7L2 revealed highly divergent β-CATENIN-dependent gene expression in different CRC cell lines. Further, β-CATENIN and TCF7L2 seemingly control predominantly non-overlapping cohorts of target genes in a given cell line. Thus, CRC cell lines may not only lose their dependence on either β-CATENIN or TCF7L2 but the close cooperation and mutual dependence of the two factors seen in healthy IECs can be broken up. While the mechanistic basis for this remains to be clarified, it is well known that β-CATENIN and TCF/LEF proteins can function separately and control expression of distinct sets of genes independently from each other in various cellular contexts including CRC cells [17],[39],[49],[51],[52]. Examples are TCF/LEF-independent transcriptional activities of β-CATENIN mediated by its interaction with Sox17 and TBX5 [17],[51], and β-CATENIN-independent functions of TCF7 and LEF1 protein in the hematopoietic system [53],[54].

The proven and eminent pathophysiological relevance of the WNT/β-CATENIN pathway in colorectal carcinogenesis make it an attractive drug target. Multiple efforts were undertaken to identify inhibitors for example of β-CATENIN/TCF7L2 transcriptional activity for therapeutic purposes [3]. However, the possibility that CRC cells quite often may not be strictly dependent on β-CATENIN or TCF7L2 and that the two factors do not necessarily cooperate in gene regulation call into question the broad suitability of the β-CATENIN/TCF7L2 complex as target for treating colorectal cancer. As an additional corollary of the observed context-dependence and divergence of β-CATENIN and TCF7L2 activities, it is virtually impossible to extrapolate from work with a small number of cell lines to colorectal cancer in general, and the significance of results from high throughput inhibitor screens which often are based on a single cell-line, is likely to be extremely limited. In this regard, though, our panel of isogenic cell line models that are either wild type or mutant for β-CATENIN and TCF/LEF family members, may serve as versatile tools to verify target specificity of drug candidates and their proposed mechanism of action.

## Methods

### Cell lines

A list with the parental wild type cell lines and the previously generated *TCF7L2*^-/-^ cell clones [10], as well as their culture conditions can be found in Suppl. Table S1.

### Genome editing

CRISPR/Cas9-mediated genome editing was performed as before [10] using sgRNAs targeting the sequences presented in Suppl. Table S12. Briefly, 1×10^6^ cells were nucleofected with 600 ng of each sgRNA expression construct, and 800 ng of Cas9-turboRFP expression vector. Single RFP^+^ cells were sorted into 96 well plates 72 h after nucleofection, expanded, and screened for deletion of the target regions by PCR with genotyping primers listed in Suppl. Table S13. For inactivation of the *TCF7* gene, HT29 *TCF7L2*^-/-^ #83.7 cells were used. These cells were derived from HT29 *TCF7L2*^-/-^ cell clone #83 [10] by lentiviral transduction with a TCF7 cDNA construct which, however, is not expressed. To obtain HCT116 triple knockout cells, HCT116 *TCF7L2*^-/-^ cell clone #3 [10] was cotransfected with the Cas9-turboRFP expression vector and four sgRNA expression vectors targeting *TCF7* and *TCF7L1*. The *CTNNB1* gene was inactivated in wild type HT29, HCT116, SW403, and RKO cells. Details about the targeting strategies and the genotypes of all cell clones generated in this study can be found in Suppl. Table S2.

### Western blotting, immunoprecipitation, and immunofluorescence experiments

For Western blotting, cells were lysed in RIPA buffer [43]. Aliquots of cell lysates with equal protein content were separated by SDS-PAGE and further processed for antigen detection as described [43]. For immunoprecipitation, whole cell lysates were prepared by lysing cells in IPN_150_ buffer [43]. To immunoprecipitate E-CADHERIN and associated proteins, aliquots of whole cell lysates with a protein content of 500 μg were combined with 500 ng anti-E-CADHERIN antibody, 30 μl of a suspension of magnetic protein G dynabeads (Life Technologies, Darmstadt, Germany), and an appropriate amount of IPN_150_ buffer to yield a final volume of 1 ml. Samples were incubated with constant agitation at 4°C overnight. Thereafter, protein G dynabeads with bound antibody/antigen complexes were collected, washed three times with IPN_150_ buffer, and finally resuspended in 1 x SDS protein loading buffer. After boiling for 5 min, proteins eluted from the beads were analyzed by Western blotting. Immunofluorescence stainings of cultured cells were carried out as described [55] using 2×10^5^ cells seeded on glass slides coated with 0.1 % gelatin. Information about antibodies and their applications is provided in Suppl. Table S14.

### β-Galactosidase staining

Cells were seeded in 6-well plates and cultivated until they had reached about 50 % confluency. β-Galactosidase staining was carried out following a published protocol with an incubation in staining solution for 16 h [56]. Pictures of the stained cells were taken using a Keyence BZ-9000 microscope.

### Measurements of population dynamics and cell cycle analyses

To compare population dynamics of WT and mutant cell lines, the cell numbers given in the corresponding figure legends were plated per well of 6-well plates and further cultivated for 72 h. Thereafter, the cells were detached and converted into single cell suspensions. Cell numbers were determined using a hemocytometer. For cell cycle analyses, 3×10^5^ cells/well were seeded in 6-well plates and incubated for 48 h. Cells were then processed for flow cytometry as described [43]. A CytoFLEX flow cytometer (Beckman Coulter, Krefeld, Germany) in combination with the FlowJo software was used for the analyses.

### Luciferase assays

To determine WNT/β-CATENIN pathway activity by reporter gene assays, 5×10^4^ cells/well were seeded in 24-well plates and transfected with 10 ng pRL-CMV (Promega, Walldorf, Germany) together with either 250 ng pSuper8xTOPFlash or 250 ng pSuper8xFOPFlash [22] using the FuGENE6® reagent (Promega). At 48 h post transfection, cells were lysed and reporter gene activities were determined as described [44]. Renilla luciferase activity was used for normalization. Where indicated, cells were treated for 40 h with 5 μM CT99021 (Selleckchem, Houston, Texas, USA) to stimulate the WNT/β-CATENIN pathway prior to measurements.

### Caspase 3 activity assay

To detect apoptotic activity, 1×10^6^ cells/well were seeded in 6-well plates and incubated for 24 h. The cells were lysed using IPN_150_-buffer with supplements (50 mM Tris/HCl, pH 7.6, 150 mM NaCl, 5 mM MgCl_2_, 0.1 % NP-40 (v/v), 10 mM NaF, 1 mM PMSF, 1 mM DTT, 0.1 mM NaVO_3_, 1 x Complete™ protease inhibitors) as previously described [43]. To measure caspase 3 activity, aliquots of whole cell lysates containing 40 ng of protein were adjusted to 20 μl volume with IPN_150_ buffer, combined with 80 μl of assay buffer (50 mM HEPES/KOH pH 7.6, 12 mM DTT) and 1 μl of DEVD-AMC substrate (final concentration 60 μm). For each cell lysate, duplicate measurements were carried out (two technical replicates). Substrate conversion was followed over a period of 30 minutes taking fluorescence readings at 1 min intervals using a TECAN Infinite F200 PRO machine (Tecan Trading AG, Männedorf, Switzerland) and excitation and emission wave lengths of 380 nm and 460 nm, respectively. For each kinetic measurement, data points were plotted, and the slope of the linear portion of the resulting curves was calculated. Values from the technical replicates were averaged, and plotted.

### Work with RNA and RNA-seq data analysis

RNA isolation, cDNA synthesis, and qRT-PCR were performed as described [43], except that a cDNA amount equivalent to 20 ng total RNA was used for qRT-PCR with oligonucleotide primers listed in Suppl. Table S13. For RNA-seq analysis, 1200 ng of total RNA were sent to Macrogen Europe for Sequencing. The library preparation was performed by Macrogen Europe using a TruSeq RNA Sample Prep Kit v2 and the according protocol (TruSeq RNA Sample Preparation v2 Guide, Part # 15026495 Rev. F). Paired-end sequencing was performed on an lllumina NovaSeq 6000 with 2×150 bp read length and 50 million read pairs per sample. For bioinformatic analysis, the galaxy.eu platform was used [57]. All reads were aligned to the human reference genome GRCh37 using RNA STAR Galaxy Version 2.7.5b [58]. For counting the reads per gene, featureCounts Galaxy Version 2.0.1 was applied [59]. DESeq2 Galaxy Version 2.11.40.6+galaxy1 was used for PCA analysis and to identify differentially expressed genes [60]. For each cell line, *CTNNB1*^-/-^ clones were compared to *CTNNB1*^+/+^ clones. Differentially expressed genes with an absolute value of log2(fold change)>1 and an adjusted p-value<0.05 were extracted (Suppl. Tables S3-S5). R was used for further data analysis and visualization of the data. For gene set enrichment analysis (GSEA), the clusterProfiler package was applied [61]. The Suppl. Tables S6-S8 contain the results of the GSEA for each cell line.

### Statistical analysis

Statistical analysis was performed using a linear mixed model (LMM) to account for random and fixed effects. Normal distribution of data was assessed using the car R package [62]. For LMM analysis, the lme4 R package was applied [63]. The ggplot2 R package [64] was used to generate box plots depicting the median, the lower and upper quartile. Whiskers represent 1.5-times the interquartile range. The ggplot2 R package [64] was used to generate bar plots depicting the mean with error bars representing the standard deviation (SD). The p-values for significant changes are represented as follows: * p < 0.05; ** p < 0.01; *** p < 0.001. For each series of experiments, the corresponding number (n) of independent biological replicates is given in the figure legends.

## Acknowledgements

The authors are grateful to the team at the Lighthouse Core Facility Freiburg, Germany, for cell sorting assistance, and all members of the Hecht laboratory for critical reading of the manuscript. The results shown here are in part based upon data generated by the TCGA Research Network: https://www.cancer.gov/tcga.

## Author contributions

J.F. and A.H. conceived and designed experiments. J.F. and K.R. performed experiments, generated, and analyzed data. J.F. carried out bioinformatic analyses of the RNA-seq data, including identification of differentially expressed genes, and gene set enrichment analyses. J.F. and A.H. created figures and tables, and wrote the manuscript. All authors critically read and commented on the final version of the manuscript.

## Data availability statement

The RNA-seq data were deposited in the Gene Expression Omnibus under the accession number GSE199835. All other data on which the results and conclusions of this study are based, are available from the corresponding author on reasonable request.

## Additional Information

**Supplementary Information** This manuscript is accompanied by supplementary material.

**Correspondence** and requests for materials should be addressed to A.H.

