## Supplementary Figures S1_S9_Supplementary Tables S1_S2_S12-S14 for "Transcriptional activity mediated by β-CATENIN and TCF/LEF family members is completely dispensable for survival of multiple human colorectal cancer cell lines"

This file contains:

Supplementary Figures S1-S9

Supplementary Tables S1, S2, S12, S13, S14

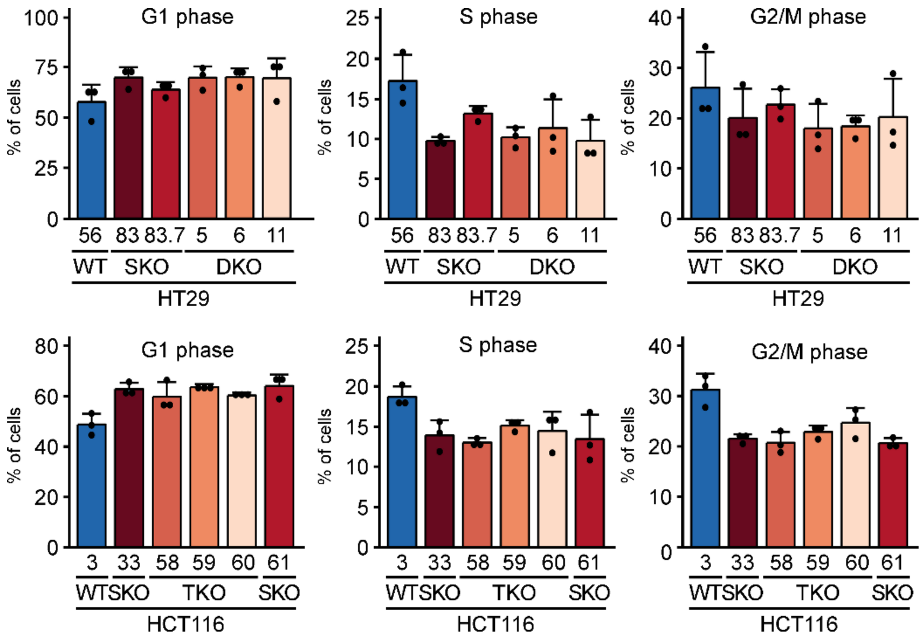


**Suppl. Fig. S1 Complete absence of TCF/LEF expression does not aggravate the proliferation defect resulting from TCF7L2-deficiency.** To examine differences in cell cycle distribution, HT29 and HCT116 cells with the genotypes indicated were stained with propidium iodide and analyzed by flow cytometry. The proportions of cells in different cell cycle phases are depicted by the bar plots. Each dot represents an individual measurement; the error bars indicate SD (n=3).

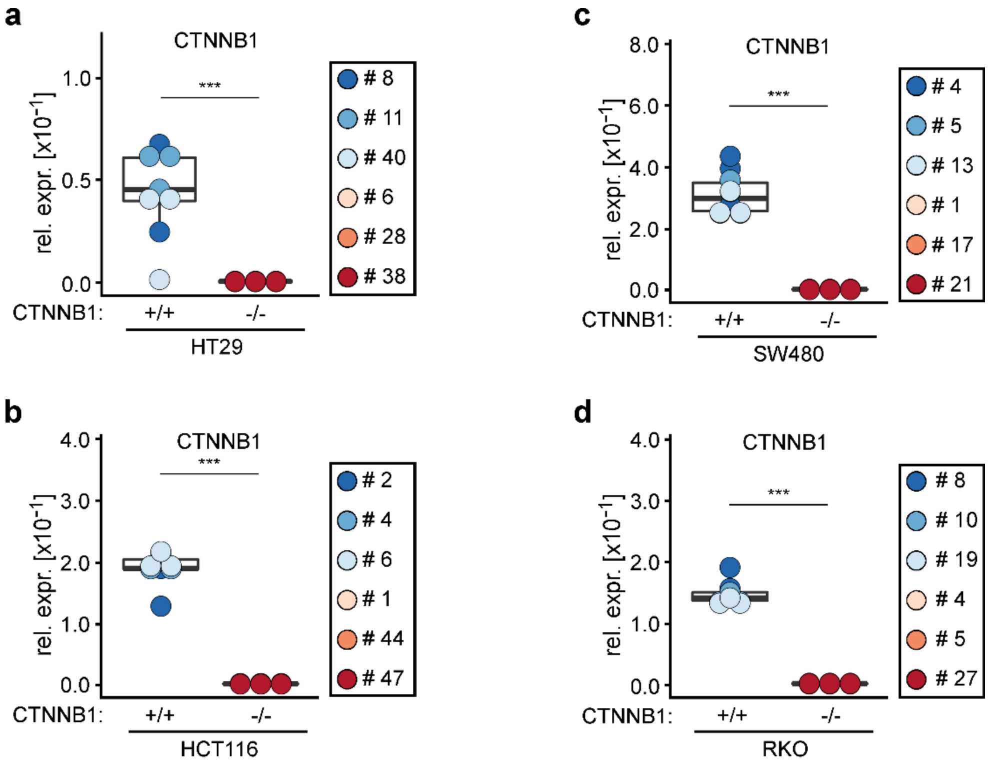


**Suppl. Fig. S2: Deletion of *CTNNB1* exon 7 results in vastly reduced RNA expression.** **a**-**d** Transcript levels of the *CTNNB1* gene were analyzed by qRT-PCR in HT29 (**a**), HCT116 (**b**), SW480 (**c**), and RKO (**d**) cell clones with biallelic wildtype and mutant *CTNNB1* genes. *CTNNB1* transcript levels were normalized to those of *GAPDH*, and are displayed as relative expression (rel. expr.). The box plots display aggregated expression data from *CTNNB1*$^{+/+}$ and *CTNNB1*$^{-/-}$ cells. Each dot represents a separate measurement; color coding identifies individual clones. For statistical analyses, LMM was performed (n=3).

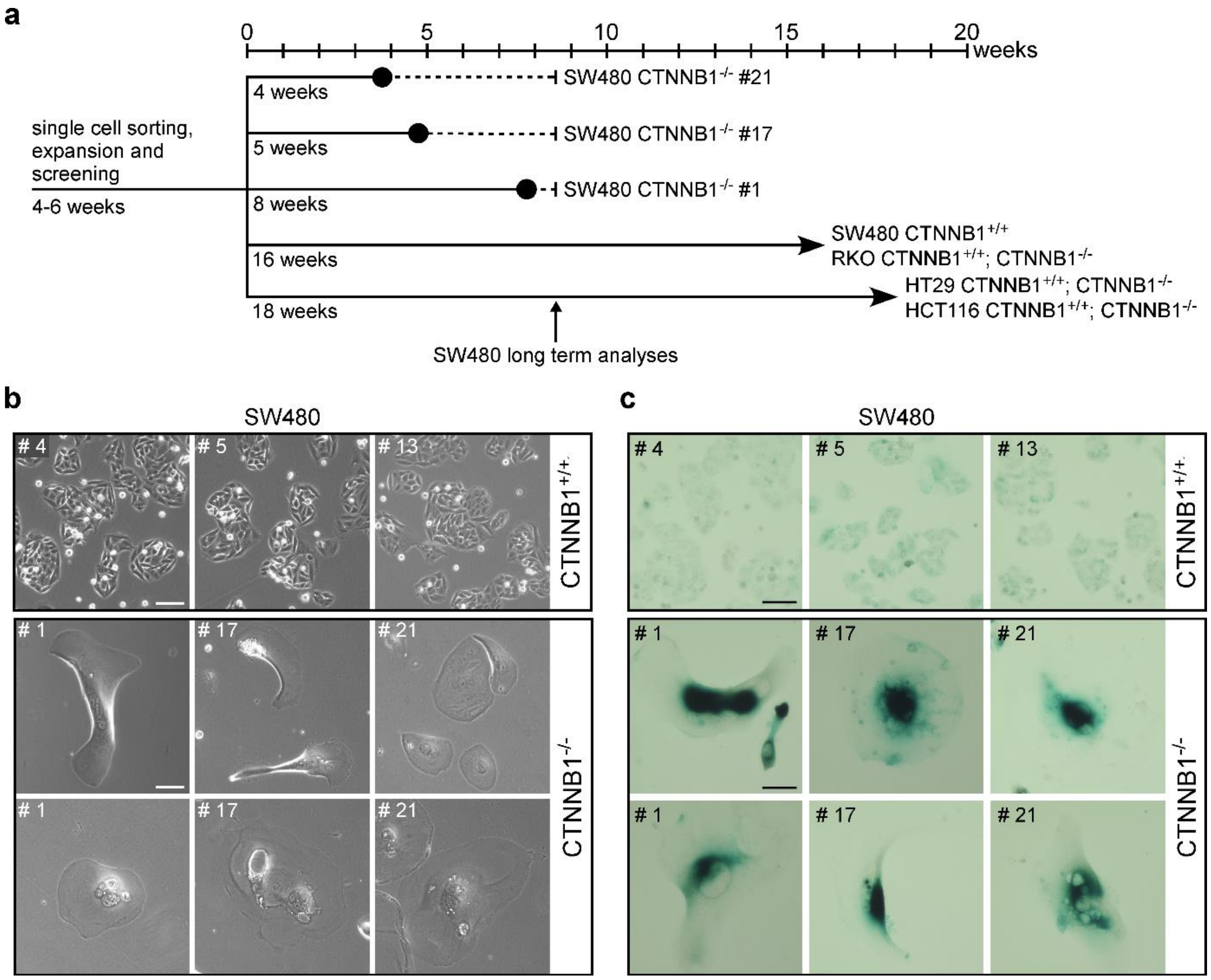


**Suppl. Fig. S3: β-CATENIN-deficient SW480 cells enter senescence after long term cell culture. a** Schematic representation of the cultivation times and growth behavior of *CTNNB1*$^{+/+}$ and *CTNNB1*$^{-/-}$ CRC cells as indicated. Large black dots represent the time points at which SW480 *CTNNB1*$^{-/-}$ cell clones stopped proliferating. Dashed lines show the additional time for which cell were kept in culture until analysis. Arrows denote that cells continue to proliferate. The vertical arrow indicates the time point at which all β-CATENIN-deficient SW480 cells were harvested and analyzed. **b** Representative micrographs showing *CTNNB1*$^{+/+}$ and *CTNNB1*$^{-/-}$ SW480 cell clones. Images from one of three independent biological replicates were taken approximately 16 weeks after expression of Cas9 and sgRNAs, and single cell-sorting. The scale bars represent 100 µm. **c** β-Galactosidase stainings of *CTNNB1*$^{+/+}$ and *CTNNB1*$^{-/-}$ SW480 cell clones performed 14 - 16 weeks after expression of Cas9 and sgRNAs, and single cell-sorting. The pictures are representative micrographs from one of three independent biological replicates. The scale bars represent 100 µm.

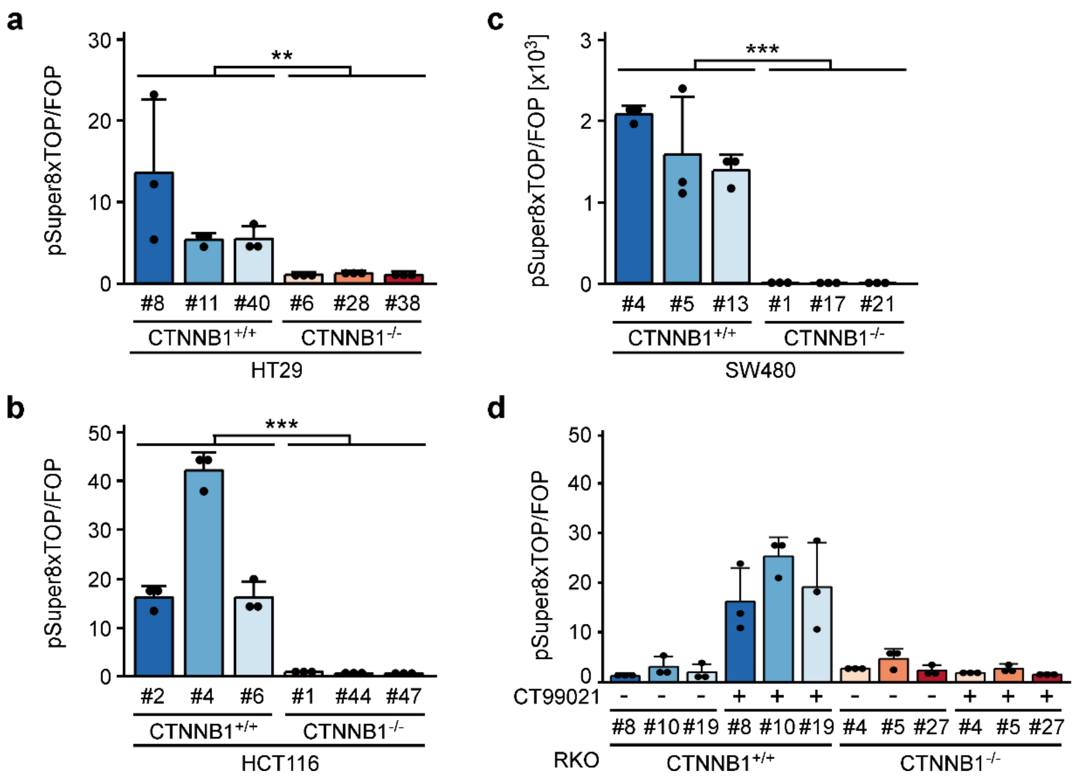


**Suppl. Fig. S4: Transcriptional activity of the Wnt/β-CATENIN pathway is abolished in β-CATENIN-deficient HT29, HCT116, SW480, and RKO cells. a-d** *CTNNB1*$^{+/+}$ and *CTNNB1*$^{-/-}$ HT29 (**a**), HCT116 (**b**), SW480 (**c**), and RKO cells (**d**) were cotransfected with expression vectors for *R. reniformis* and firefly luciferase reporter genes with wild type (pSuper8xTOPFlash) or mutant (pSuper8xFOPFlash) TCF/LEF binding sites in their promoter elements. RKO cells were additionally treated with CT99021 to stimulate Wnt pathway activity. Luciferase activities were determined 48 h post transfection. *R. reniformis* luciferase activities were used for normalization of firefly luciferase activities. The bar graphs show the ratios of normalized firefly luciferase activities from cells transfected with pSuper8xTOPFlash and with pSuper8xFOPFlash. Each dot represents an individual measurement. Error bars indicate SD (n=3). For statistical analysis LMM was performed.

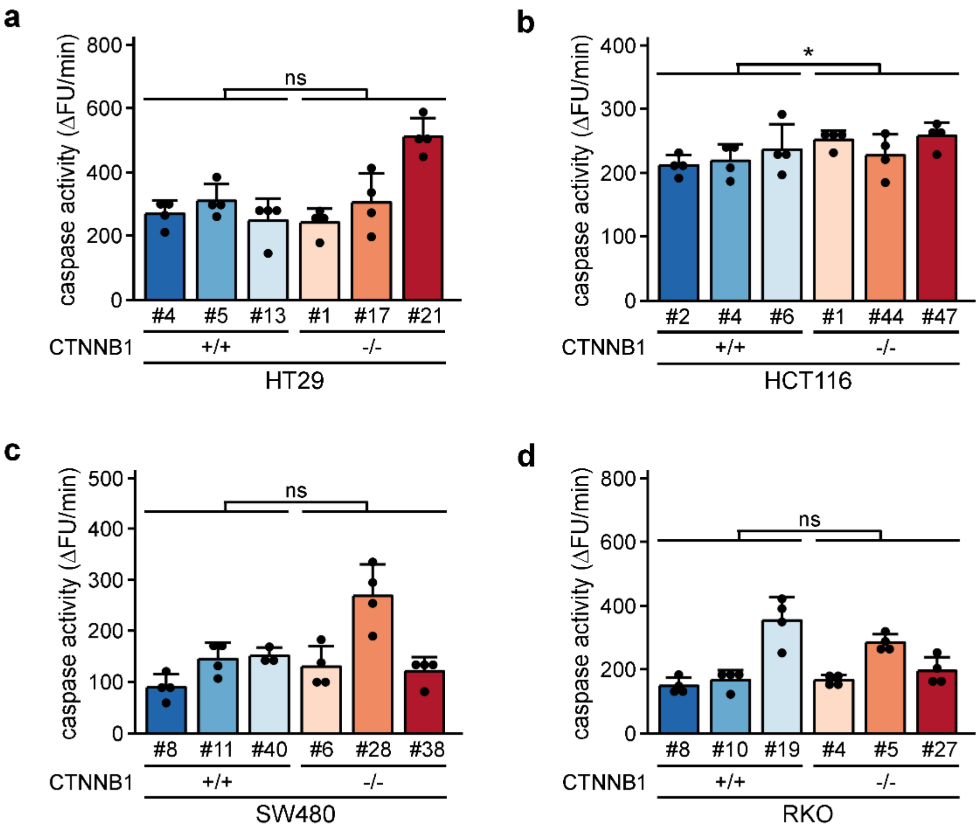


**Suppl. Fig. S5: β-CATENIN-deficiency does not lead to increased apoptosis in HT29, HCT116, SW480, and RKO cells. a-d** *CTNNB1*$^{+/+}$ and *CTNNB1*$^{-/-}$ HT29 (**a**), HCT116 (**b**), SW480 (**c**), and RKO (**d**) cells were seeded and incubated for 24 h. Upon cell lysis and preparation of whole cell extracts, caspase 3 activity was determined by a fluorimetric assay and kinetic measurements. The resulting changes in fluorescence units (ΔFU) per min are displayed in the bar plots. Each dot represents an individual measurement. Error bars indicate SD (n=3). For statistical analysis LMM was performed. ns: not significant.

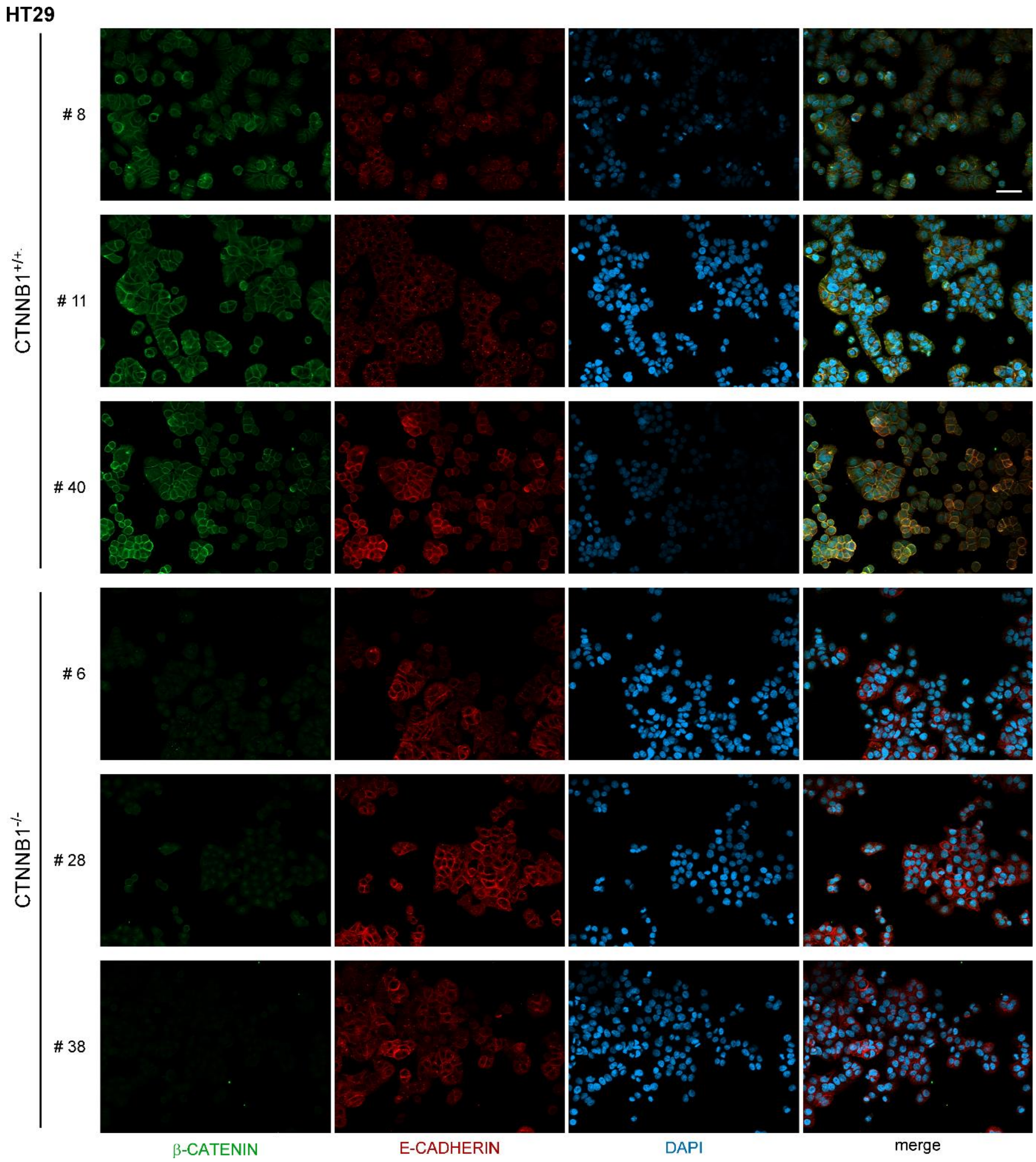


**Suppl. Fig. S6 Membrane localization of E-CADHERIN is maintained in β-CATENIN-deficient HT29 cells.** *CTNNB1*$^{+/+}$ and *CTNNB1*$^{-/-}$ HT29 cell clones were seeded on gelatin-coated glass slides and incubated for 24 h. After fixation, E-CADHERIN and β-CATENIN were visualized by immunofluorescence stainings with specific antibodies. Nuclei were counterstained with DAPI (blue color). The pictures are representative micrographs from one of three independent biological replicates. The scale bar represents 50 µm.

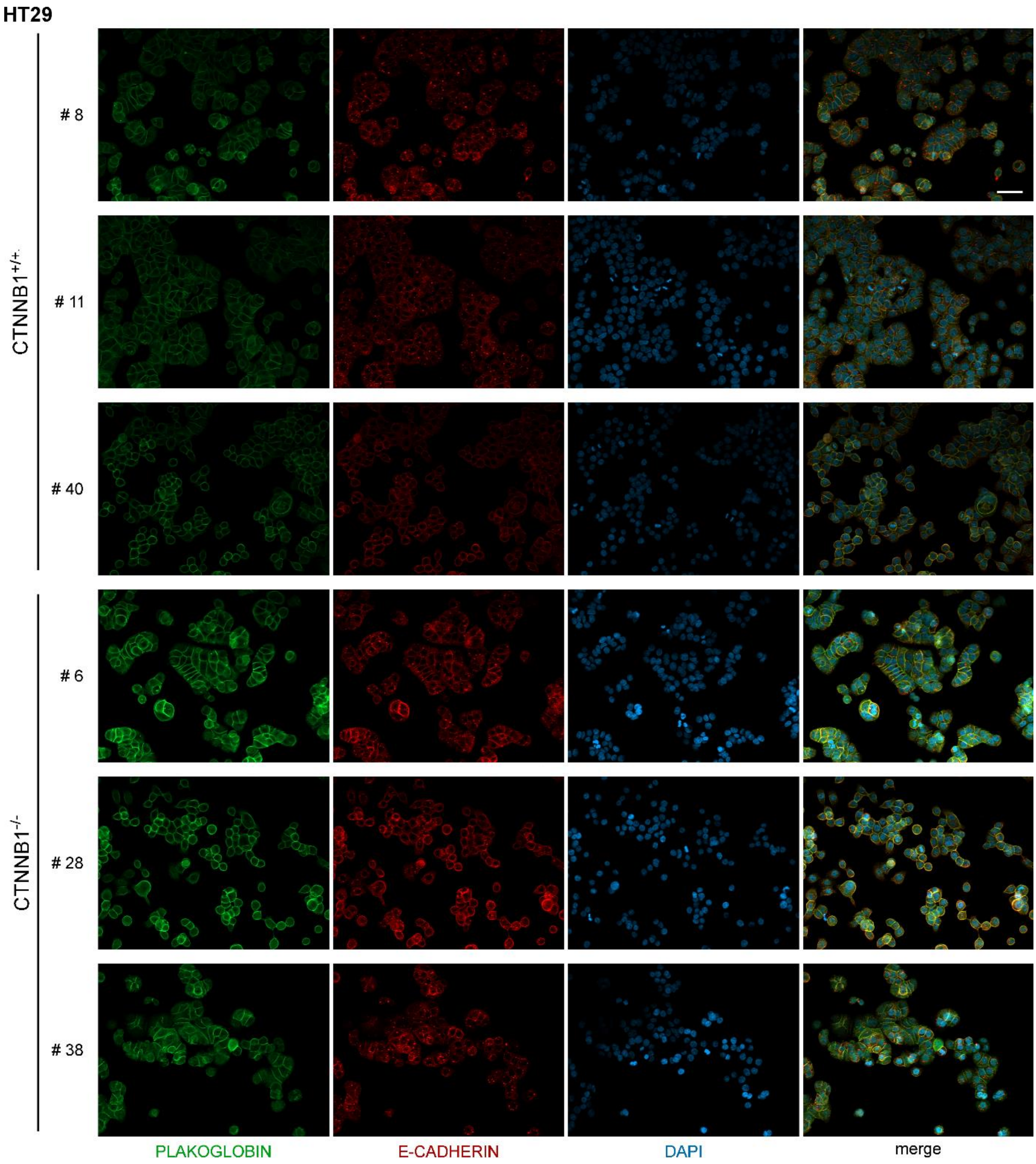


**Suppl. Fig. S7: Plakoglobin localizes to cell-cell interfaces in *CTNNB1*$^{+/+}$ and *CTNNB1*$^{-/-}$ HT29 cells.** *CTNNB1*$^{+/+}$ and *CTNNB1*$^{-/-}$ HT29 cell clones were seeded on gelatin-coated glass slides and incubated for 24 h. After fixation, E-CADHERIN and PLAKOGLOBIN were visualized by immunofluorescence stainings with specific antibodies. Nuclei were counterstained with DAPI (blue color). The pictures are representative micrographs from one of three independent biological replicates. Scale bar represents 50 µm.

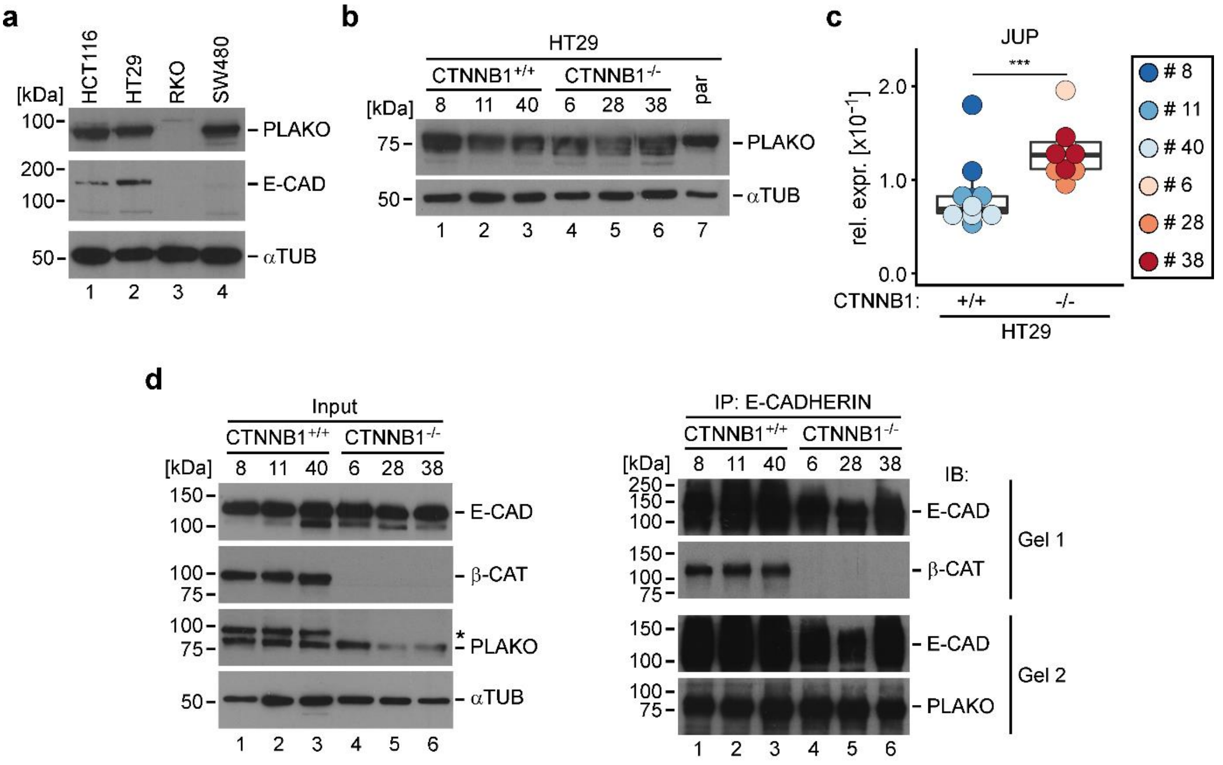


**Suppl. Fig. S8: PLAKOGLOBIN is expressed in CRC cell lines and interacts with E-CADHERIN. a** Western blot experiment to detect expression of E-CADHERIN (E-CAD) and PLAKOGLOBIN (PLAKO) in the CRC cancer cell lines shown. α-TUBULIN (α-TUB) was used to demonstrate equal loading. Molecular weights are given in kDa. **b** Western blot experiment to detect expression of PLAKOGLOBIN in *CTNNB1*$^{+/+}$ and *CTNNB1*$^{-/-}$ HT29 cell clones. For comparison, cell lysates from parental (par) HT29 cells were analyzed in parallel. α-TUBULIN (αTUB) was used to demonstrate equal loading. Molecular weights are given in kDa. **c** Transcript levels of the *JUP* gene (coding for PLAKOGLOBIN) were analyzed by qRT-PCR in *CTNNB1*$^{+/+}$ and *CTNNB1*$^{-/-}$ HT29 cell clones. *JUP* transcript levels were normalized to those of *GAPDH*, and are displayed as relative expression (rel. expr.). The box plots display aggregated expression data whereby each dot represents a separate measurement, while dot color identifies individual clones. For statistical analysis, LMM was performed (n=3). **d** Whole cell lysates from *CTNNB1*$^{+/+}$ and *CTNNB1*$^{-/-}$ HT29 cell clones were prepared and subjected to immunoprecipitation (IP) with anti-E-CADHERIN antibodies. Immunoprecipitates were split into two halves each of which was loaded onto a separate SDS-polyacrylamide gel and analyzed by immunoblotting (IB) with antibodies directed against β-CATENIN (β-CAT), E-CADHERIN, and PLAKOGLOBIN as depicted. Aliquots of the input material were analyzed in parallel. β-CATENIN and PLAKOGLOBIN in the input were detected by sequential probing of the same membrane strip which explains the reappearance of the β-CATENIN signal (marked by an asterisk) during detection of PLAKOGLOBIN. α-TUBULIN was used to demonstrate equal loading. Molecular weights are given in kDa. **a**, **b**, **d** Representative results from one of three independent biological replicates are shown. Full size/uncropped versions of the Western blot images are shown in Suppl. Fig. S9.

### Suppl. Fig. S9

#### Related to Figure 1b:

Gel 1

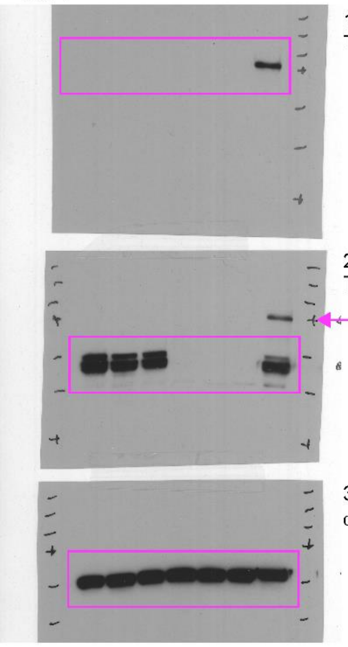


Gel 2

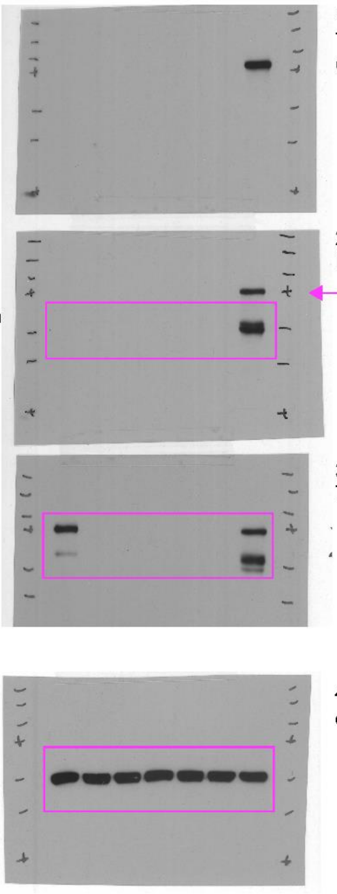


#### Related to Figure 1c:

Gel 1

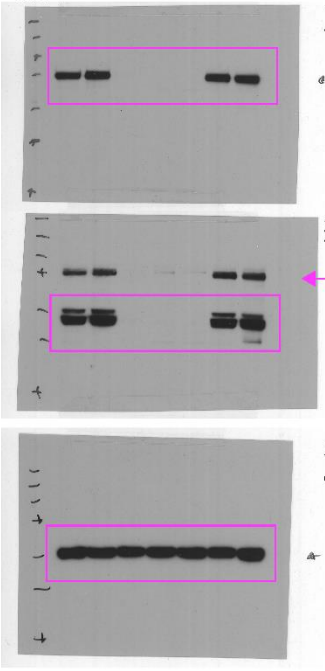


Gel 2

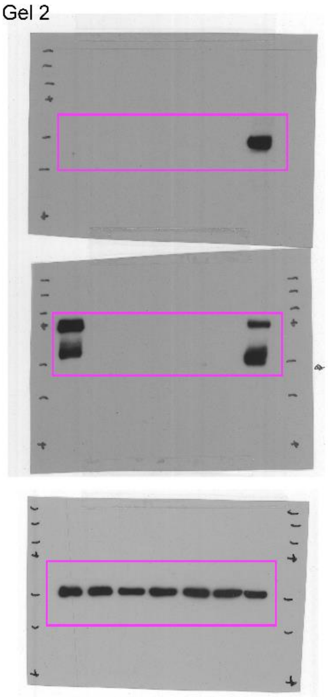


**Suppl. Fig. S9 continued →**

**Suppl. Fig. S9 continued:**

**Related to Figure 3b:**

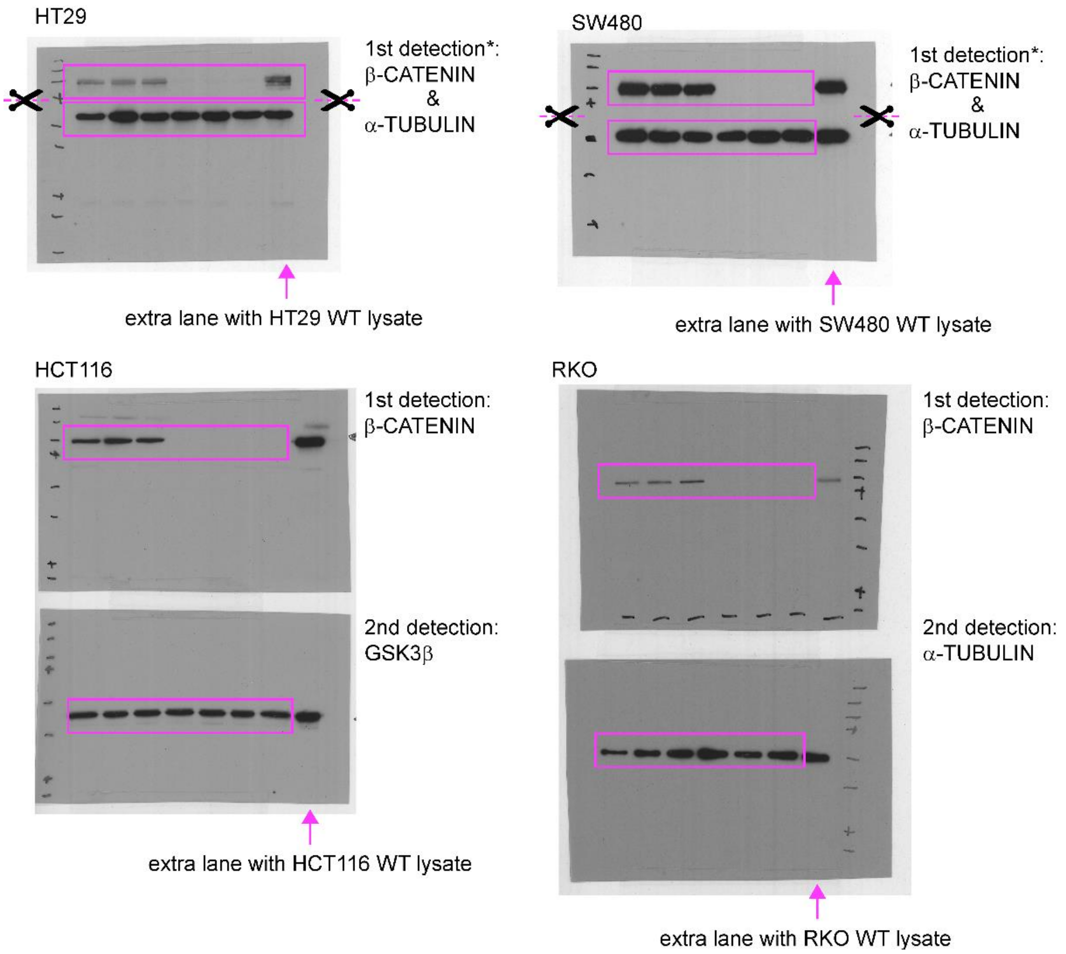


*: after the gel run and transfer to nitrocellulose, the membrane was cut horizontally below the 75 kDa marker (4th from top, marked with a cross). The top and bottom parts of the membrane were then processed separately for detection of β-CATENIN and α-TUBULIN. However, for signal detection, the two pieces of membrane were reassembled.

**Suppl. Fig. S9 continued →**

### Suppl. Fig. S9 continued

#### Related to Suppl. Fig. S8a:

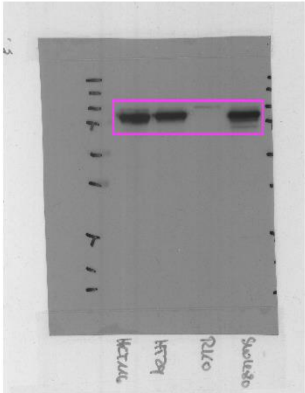

1st detection:
PLAKOGLOBIN

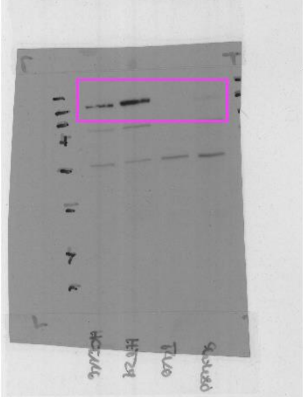

2nd detection:
E-CADHERIN

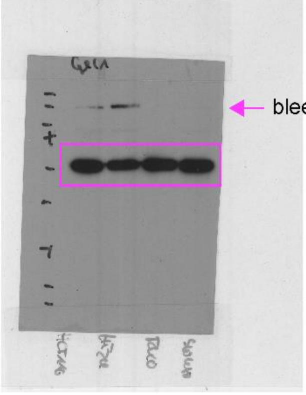

3rd detection:
α-TUBULIN
← bleed through from 2nd detection

#### Related to Suppl. Fig. S8b:

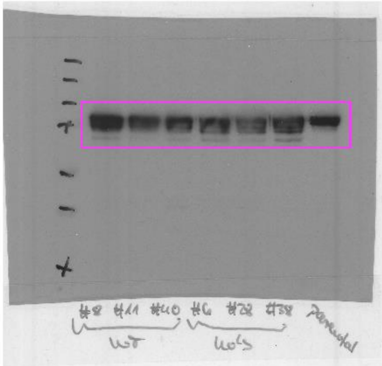

1st detection:
PLAKOGLOBIN

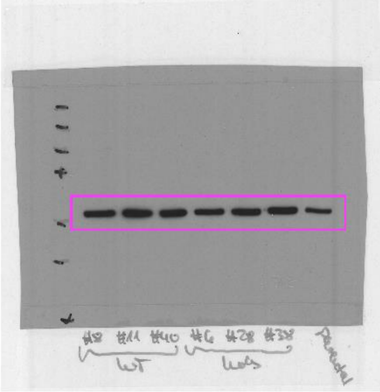

2nd detection:
α-TUBULIN

**Suppl. Fig. S9 continued →**

**Suppl. Fig. S9 continued**

**Related to Suppl. Fig. S8d, Input:**

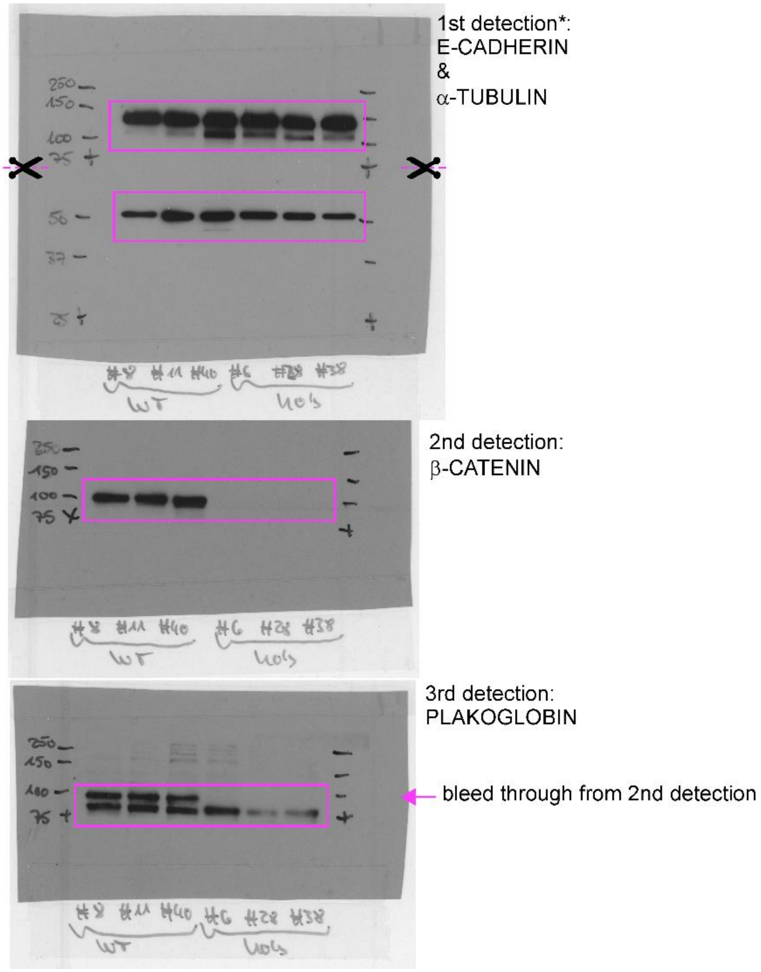


*: after the gel run and transfer to nitrocellulose, the membrane was cut horizontally below the 75 kDa marker (4th from top, marked with a cross). The top and bottom parts of the membrane were then processed separately for detection of E-CADHERIN and α-TUBULIN. However, for signal detection, the two pieces of membrane were reassembled.

**Suppl. Fig. S9 continued →**

### Suppl. Fig. S9 continued

**Related to Suppl. Fig. S8d, coIP:**

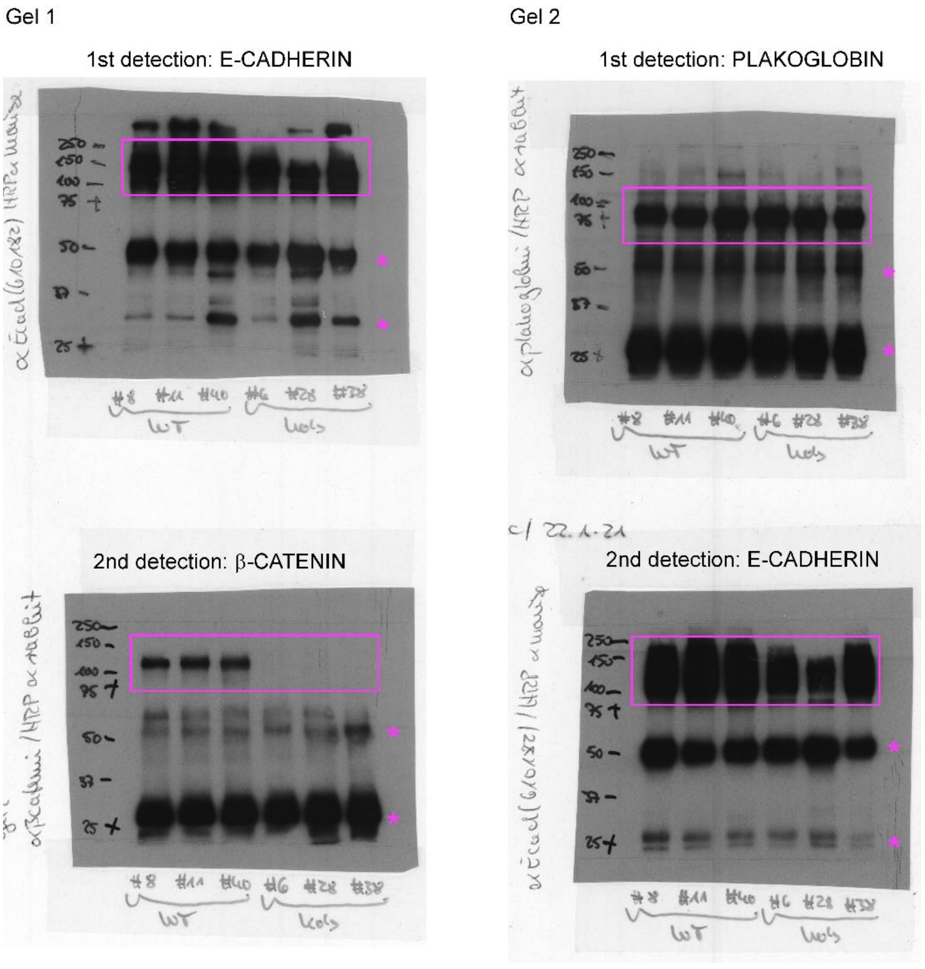


*: IgG heavy and light chains

**Suppl. Fig. S9: Full size images of Western blots shown Figs. 1 and 3 of the main manuscript and Suppl. Fig. S8 of the Supplementary Information.** Pink frames highlight areas used for display in the final versions of the figures. When membranes were sequentially probed, the order of detection is given and bleed-through signals reappearing during subsequent rounds of the detection, are marked.

### Supplementary Table S1: Cell lines used in this study

| Cell line*,# | Source | Culture conditions |
|---|---|---|
| HCT116 | Max-Planck-Institute for Immunology and Epigenetics (Freiburg, Germany) | DMEM with 4.5 g/l glucose, stabilized glutamine, sodium pyruvate, and 3.7 g/l $NaHCO_3$, supplemented with:<br>10 % (v/v) fetal calf serum<br>10 mM HEPES<br>1 % (v/v) MEM non-essential amino acids solution<br>1 % (v/v) penicillin/streptomycin<br><br>grown at 37°C and 5 % $CO_2$. |
| HCT116 $TCF7L2^{+/+}$ clone 3 | derived from HCT116[§] | |
| HCT116 $TCF7L2^{-/-}$ clone 33 | derived from HCT116[§] | |
| HT29 | German Cancer Research Center Cell Line Service (Heidelberg, Germany) | |
| HT29 $TCF7L2^{+/+}$ clone 56 | derived from HT29[§] | |
| HT29 $TCF7L2^{-/-}$ clone 83 | derived from HT29[§] | |
| LoVo | CLS Cell Lines Service GmbH (Eppelheim, Germany) | |
| LS174T | CLS Cell Lines Service GmbH (Eppelheim, Germany) | |
| RKO | Institute of Molecular Medicine and Cell Research (Freiburg, Germany) | |
| SW480 | Max-Planck-Institute for Immunology and Epigenetics (Freiburg, Germany) | |

* Cell line identity was determined by SNP-profiling at Multiplexion Inc. (Friedrichshafen, Germany).

[#] Cell lines were routinely tested for mycoplasma contamination using the Myco sensor PCR assay kit from Stratagene (San Diego, CA, USA).

[§] generation of these cell lines was described before (Wenzel et al., 2020).

### Supplementary Table S2: Summary of genome editing strategies and genotypes of cell clones generated in this study

| Derived from HT29 *TCF7L2*$^{-/-}$ clone #83.7* (Wenzel et al., 2020) | | | |
|---|---|---|---|
| targeted gene: *TCF7* | | | |
| clone ID | sgRNAs used | genotype | allele 1/2 |
| #5 | sgTCF7-1<br>sgTCF7-2 | KO | ∆ ENSE00003616545 |
| #6 | sgTCF7-1<br>sgTCF7-2 | KO | ∆ ENSE00003616545 |
| #11 | sgTCF7-1<br>sgTCF7-2 | KO | ∆ ENSE00003616545 |

* For inactivation of *TCF7*, HT29 *TCF7L2*$^{-/-}$ #83.7 cells were used. These cells are derived from HT29 *TCF7L2*$^{-/-}$ cell clone #83 by lentiviral transduction with a TCF7 cDNA construct which, however, is not expressed.

| Derived from HCT116 *TCF7L2*$^{-/-}$ clone #3 (Wenzel et al., 2020) | | | | | | |
|---|---|---|---|---|---|---|
| clone ID | targeted gene: *TCF7* | | | targeted gene: *TCF7L1* | | |
| | sgRNAs used | genotype | allele 1/2 | sgRNAs used | genotype | allele 1/2 |
| #61 | sgTCF7-1<br>sgTCF7-2 | WT | WT | sgTCF7L1-1<br>sgTCF7L1-2 | WT | WT |
| #58 | sgTCF7-1<br>sgTCF7-2 | KO | ∆ ENSE00003616545 | sgTCF7L1-1<br>sgTCF7L1-2 | KO | 209 bp deletion; nucleotides 28-128 of ENSE00000963602[a] |
| #59 | sgTCF7-1<br>sgTCF7-2 | KO | ∆ ENSE00003616545 | sgTCF7L1-1<br>sgTCF7L1-2 | KO | 272 bp deletion; nucleotides 1-78 of ENSE00000963602[a] |
| #60 | sgTCF7-1<br>sgTCF7-2 | KO | ∆ ENSE00003616545 | sgTCF7L1-1<br>sgTCF7L1-2 | KO | 209 bp deletion; nucleotides 28-128 of ENSE00000963602[a] |

| derived from HCT116 cells | | | |
|---|---|---|---|
| targeted gene: *CTNNB1* | | | |
| clone ID | sgRNAs used | genotype | allele 1/2 |
| #2 | sgCTNNB1-1<br>sgCTNNB1-2 | WT | WT |
| #4 | sgCTNNB1-1<br>sgCTNNB1-2 | WT | WT |
| #6 | sgCTNNB1-1<br>sgCTNNB1-2 | WT | WT |
| #1 | sgCTNNB1-1<br>sgCTNNB1-2 | KO | 258 bp deletion; nucleotides 17-145 of ENSE00001643204[b] |
| #44 | sgCTNNB1-1<br>sgCTNNB1-2 | KO | 258 bp deletion; nucleotides 17-145 of ENSE00001643204[b] |
| #47 | sgCTNNB1-1<br>sgCTNNB1-2 | KO | 258 bp deletion; nucleotides 17-145 of ENSE00001643204[b] |

continued →

| derived from HT29 cells | | | |
|---|---|---|---|
| **targeted gene: *CTNNB1*** | | | |
| **clone ID** | **sgRNAs used** | **genotype** | **allele 1/2** |
| #8 | sgCTNNB1-1<br>sgCTNNB1-2 | WT | WT |
| #11 | sgCTNNB1-1<br>sgCTNNB1-2 | WT | WT |
| #40 | sgCTNNB1-1<br>sgCTNNB1-2 | WT | WT |
| #6 | sgCTNNB1-1<br>sgCTNNB1-2 | KO | 258 bp deletion; nucleotides 17-145 of ENSE00001643204[b] |
| #28 | sgCTNNB1-1<br>sgCTNNB1-2 | KO | 258 bp deletion; nucleotides 17-145 of ENSE00001643204[b] |
| #38 | sgCTNNB1-1<br>sgCTNNB1-2 | KO | 258 bp deletion of nucleotides 17-145 of ENSE00001643204[b] |

| derived from SW480 cells | | | |
|---|---|---|---|
| **targeted gene: *CTNNB1*** | | | |
| **clone ID** | **sgRNAs used** | **genotype** | **allele 1** |
| #4 | sgCTNNB1-1<br>sgCTNNB1-2 | WT | WT |
| #5 | sgCTNNB1-1<br>sgCTNNB1-2 | WT | WT |
| #13 | sgCTNNB1-1<br>sgCTNNB1-2 | WT | WT |
| #1 | sgCTNNB1-1<br>sgCTNNB1-2 | KO | 258 bp deletion; nucleotides 17-145 of ENSE00001643204[b] |
| #17 | sgCTNNB1-1<br>sgCTNNB1-2 | KO | 258 bp deletion; nucleotides 17-145 of ENSE00001643204[b] |
| #21 | sgCTNNB1-1<br>sgCTNNB1-2 | KO | 258 bp deletion; nucleotides 17-145 of ENSE00001643204[b] |

| derived from RKO cells | | | |
|---|---|---|---|
| **targeted gene: *CTNNB1*** | | | |
| **clone ID** | **sgRNAs used** | **genotype** | **allele 1/2** |
| #8 | sgCTNNB1-1<br>sgCTNNB1-2 | WT | WT |
| #10 | sgCTNNB1-1<br>sgCTNNB1-2 | WT | WT |
| #19 | sgCTNNB1-1<br>sgCTNNB1-2 | WT | WT |
| #4 | sgCTNNB1-1<br>sgCTNNB1-2 | KO | 258 bp deletion; nucleotides 17-145 of ENSE00001643204[b] |
| #5 | sgCTNNB1-1<br>sgCTNNB1-2 | KO | 258 bp deletion; nucleotides 17-145 of ENSE00001643204[b] |
| #27 | sgCTNNB1-1<br>sgCTNNB1-2 | KO | 258 bp deletion; nucleotides 17-145 of ENSE00001643204[b] |

[a] coordinates refer to the 5´-end of ENSE00000963602
[b] coordinates refer to the 5´-end of ENSE00001643204

**Supplementary Table S12: List of sgRNA target sequences (without PAM)**

| Gene | sgRNA | Location | Sequence (5` - 3`) |
|---|---|---|---|
| CTNNB1 | sgCTNNB1-1 | exon 7 | CTCATCATACTGGCTAGTGG |
| CTNNB1 | sgCTNNB1-2 | intron 7 | GGTACTCTGAATGTAAATCT |
| TCF7 | sgTCF7-1 | intron 3 | GTGAGTGTGGCGAGTCCTGA |
| TCF7 | sgTCF7-2 | intron 4 | CCTGGGGCTGTGCAAACTAA |
| TCF7L1 | sgTCF7L1-1 | exon 3 | TTAAAGAACGCGCTGTCCTG |
| TCF7L1 | sgTCF7L1-1 | intron 3 | CTGCTTGGGATCGGCGCAGA |

**Supplementary Table S13: List of oligonucleotide primers used in this study**

| Primers for genotyping | | |
|---|---|---|
| **Gene** | **Forward primer (5` - 3`)** | **Reverse primer (5` - 3`)** |
| CTNNB1 | GGACAAGTTGGATAGGGCCC | GCACACGAAACCCCTGTGA |
| TCF7 | GCAAAGTCTTGGGGGCTAGT | GGGTCACCCATGGGATTTAGG |
| TCF7L1 | GCTCACCCGCTCTTGCCTTTGT | GAGGACAACGTCGCCAACCCAG |
| **Primers for qRT-PCR** | | |
| **Gene** | **Forward primer (5` - 3`)** | **Reverse primer (5` - 3`)** |
| ASCL2 | TGACCTGGGGCGTAATAAAG | ACACAGGCTTCTCCCTAGCA |
| AXIN2 | TGCTTTCGTGGAAATGACAG | AGGTGTGTGGAGGAAAGGTG |
| CTNNB1 | ACTGGCAGCAACAGTCTTAC | GTGGCAAGTTCTGCATCATC |
| GAPDH | ACCACAGTCCATGCCATCACT | GTCCACCACCCTGTTGCTGTA |
| LEF1 | ACAGATCACCCCACCTCTTG | TGAGGCTTCACGTGCATTAG |
| MYC | AAGAGGACTTGTTGCGGAAA | CTCAGCCAAGGTTGTGAGGT |
| RNF43 | CTGCTACCAGAAACCCCAGG | CTGCGGTGTCAGAACTCCAT |
| TCF7 | AGCCAAGGTCATTGCAGAGT | GTGGTGGATTCTTGGTGCTT |
| TCF7L1 | GGGTACCCCTTCCTGATGAT | GATGGTGACCTCGTGTCCTT |
| TCF7L2 | AGAAAAGAAGAAGCCCCACA | CGGGCCAGCTCGTAGTATT |
| TERT | CTACGGCGACATGGAGAACA | AGAGATGACGCGCAGGAAAA |

### Supplementary Table S14: List of antibodies used in this study

| Antigen | Species of origin | Dilution | | | Supplier (catalogue no.) |
|---|---|---|---|---|---|
| | | WB* | IP§ | IF# | |
| α-TUBULIN | mouse | 1:10 000 | - | - | Sigma-Aldrich (T9026) |
| β-CATENIN | rabbit | 1:1000 | - | 1:200 | Cell Signaling Technology (D10A8) |
| E-CADHERIN | mouse | - | 1:500 | 1:200 | BD Biosciences (610182) |
| E-CADHERIN | mouse | 1:1000 | - | - | BD Biosciences (610404) |
| LEF1 | rabbit | 1:1000 | - | - | Cell Signaling Technology (C12A5) |
| PLAKOGLOBIN | rabbit | 1:1000 | - | 1:200 | Thermo Fisher Scientific (A303-718A) |
| TCF7 | rabbit | 1:1000 | - | - | Cell Signaling Technology (C63D9) |
| TCF7L1 | rabbit | 1:1000 | - | - | Cell Signaling Technology (D15G11) |
| TCF7L2 | rabbit | 1:1000 | - | - | Cell Signaling Technology (C9B9) |

*WB: Western blotting

§IP: Immunoprecipitation

#IF: Immunofluorescence
